# BRAKER3: Fully automated genome annotation using RNA-seq and protein evidence with GeneMark-ETP, AUGUSTUS and TSEBRA

**DOI:** 10.1101/2023.06.10.544449

**Authors:** Lars Gabriel, Tomáš Brůna, Katharina J. Hoff, Matthis Ebel, Alexandre Lomsadze, Mark Borodovsky, Mario Stanke

## Abstract

Gene prediction has remained an active area of bioinformatics research for a long time. Still, gene prediction in large eukaryotic genomes presents a challenge that must be addressed by new algorithms. The amount and significance of the evidence available from transcriptomes and proteomes vary across genomes, between genes and even along a single gene. User-friendly and accurate annotation pipelines that can cope with such data heterogeneity are needed. The previously developed annotation pipelines BRAKER1 and BRAKER2 use RNA-seq or protein data, respectively, but not both. A further significant performance improvement was made by the recently released GeneMark-ETP integrating all three data types.

We here present the BRAKER3 pipeline that builds on GeneMark-ETP and AUGUSTUS and further improves accuracy using the TSEBRA combiner. BRAKER3 annotates protein-coding genes in eukaryotic genomes using both short-read RNA-seq and a large protein database, along with statistical models learned iteratively and specifically for the target genome. We benchmarked the new pipeline on genomes of 11 species under assumed level of relatedness of the target species proteome to available proteomes. BRAKER3 outperformed BRAKER1 and BRAKER2. The average transcript-level F1-score was increased by *∼*20 percentage points on average, while the difference was most pronounced for species with large and complex genomes. BRAKER3 also outperformed other existing tools, MAKER2, Funannotate and FINDER. The code of BRAKER3 is available on GitHub and as a ready-to-run Docker container for execution with Docker or Singularity. Overall, BRAKER3 is an accurate, easy-to-use tool for eukaryotic genome annotation.

## Introduction

New eukaryotic genomes are being sequenced at increasing rates. However, the pace of genome annotation, which establishes links between genomic sequence and biological function, is lagging behind. For example, in April 2023, 49% of the eukaryotic species with assemblies in GenBank had no annotation in GenBank. Undertakings such as the Earth BioGenome Project (https://www.earthbiogenome.org), which aims to annotate *∼*1.5 million eukaryotic species, further require that the annotation pipeline is highly automated and reliable and ideally no manual work for each species is required when genome assembly and RNA-seq are given.

Further, species which have an annotation also require re-annotation as assemblies improve or the available extrinsic evidence increases substantially (https://www.ncbi.nlm.nih.gov/genome/annotation_euk/). This demand further increases the importance of the availability of fast and accurate genome annotation tools.

Current state-of-the-art annotation pipelines integrate extrinsic and intrinsic evidence. Extrinsic evidence is extracted from transcripts and cross-species homologous proteins. RNA-seq reads offer direct evidence on introns and, if assembled, on a gene structure. Protein sequences from related genomes can be used to identify regions of a genome that encode proteins with similar sequences to known proteins. Due to the sequence divergence between informant and target gene, this evidence may be less reliable and less precise than the one from (native) RNA-seq alignments. The availability of extrinsic evidence is increasing rapidly. Second-generation sequencing technology has become cheap and RNA-seq often accompanies genome sequencing, as reported by Wetterstrand in 2021 (https://www.genome.gov/about-genomics/fact-sheets/Sequencing-Human-Genome-cost). To give an example for protein database growth, OrthoDB’s latest release (v11) includes more than 50% additional eukaryotic species compared to its previous version (Kuznetsov et al., 2023).

Despite the importance of extrinsic evidence, it may cover only some parts of a gene, leaving other parts without evidence. Traditional *ab initio* gene prediction methods rely on computational predictions by statistical models using genome sequence data alone, for example AUGUSTUS (Stanke et al., 2006) and GeneMark-ES (Lomsadze et al., 2005). However, the *ab initio* models are prone to errors when used alone. Therefore, more precise gene predictions are made when predictions based on statistical models are corrected by extrinsic evidence (Stanke et al., 2008; Lomsadze et al., 2014; Brůna et al., 2020).

Earlier developed BRAKER1 (Hoff et al., 2016) and BRAKER2 (Brůna et al., 2021) combined GeneMark and AUGUSTUS to utilize, respectively, a single source of extrinsic evidence, either RNA-seq short reads or homologous proteins. The use of both extrinsic evidence sources together has a clear potential for more accurate gene structure prediction. Therefore, we developed a combiner tool TSEBRA (Gabriel et al., 2021). It selects transcripts from BRAKER1 and BRAKER2 annotations, considering thereby the joint extrinsic evidence and, therefore, generates a prediction based on both RNA-seq and protein evidence, thus improving the F1-scores.

A more integrated approach is the GeneMark-ETP pipeline (Brůna et al., 2023b), which integrates both sources of extrinsic evidence in a new workflow that outperforms all previously mentioned methods, particularly in species with large and complex genomes. Critical to its improvement is a novel approach to generate a training set that has a precision from genes predicted in assembled transcripts and supported by protein evidence. The method also benefited from the GC-content-specific model training, and estimating species-specific repeat penalties.

These many advancements and the steady increase in popularity of the previous BRAKER tools motivated us to develop a new version of the BRAKER pipeline that can utilize both transcript and protein homology extrinsic evidence by incorporating GeneMark-ETP, AUGUSTUS, and TSEBRA into a novel workflow.

Similar tools that use RNA-seq and protein data are MAKER2 (Holt and Yandell, 2011), FINDER (Banerjee et al., 2021) and Funannotate (https://github.com/nextgenusfs/funannotate). MAKER2 aligns assembled RNA-seq data and proteins to the genome and can run and integrate SNAP (Korf, 2004), GeneMark and AUGUSTUS predictions. Although MAKER2 can provide training sets for SNAP and AUGUSTUS, it does not train the *ab initio* models automatically. Also, the self-training of GeneMark.hmm models (Lomsadze et al., 2005) has to be done outside of MAKER2. FINDER follows an approach similar to BRAKER3. It uses RNA-seq assemblies with predicted open reading frames, in conjunction with BRAKER1 and homologous protein predictions. The Funannotate pipeline, which was not described in an article, was initially designed as a pipeline for analyzing fungal genomes; however, it has since been further developed to support the annotation of larger genomes as well (Palmer, 2017).

In computational experiments with genomes of 11 species we have accessed and compared performances of BRAKER1, BRAKER2, TSEBRA, GeneMark-ETP and BRAKER3. Also, we have conducted several experiments to access and compare performances of BRAKER3 with FINDER, Funannotate and MAKER2. We have demonstrated that BRAKER3 consistently outperformed the other gene finding tools.

## Methods

### BRAKER3

BRAKER3 is the latest genome annotation pipeline that continues the BRAKER family. It requires three types of inputs: the genome sequence to annotate, a list of short-read RNA-seq datasets and a protein database file. The protein database is a `FASTA` file with proteins from the broad clade of the target genome in question, e.g., a subset from the partitioning of OrthoDB that we provide (see Suppl. Methods). To specify the RNA-seq input there are three options: as `BAM`-files of aligned reads, as raw reads in `FASTQ`-files, or as SRA (Leinonen et al. (2010)) library IDs.

BRAKER3 runs the GeneMark-ETP pipeline which performs the steps that are outlined next and described in detail in Brůna et al. (2023b). First, transcript sequences are assembled with StringTie2 (Kovaka et al., 2019) from the short RNA-seq reads aligned to the genome by HISAT2 (Kim et al., 2019). The assembled transcripts are then analyzed by GeneMarkS-T (Tang et al., 2015) to predict the protein-coding genes. The predicted proteins are searched against the protein database, and GeneMark-ETP uses the resulting similarity scores to identify *high confidence* gene structures. Then, the parameters of GeneMark.hmm are trained on the high confidence genes, and it predicts genes in the *intermediate fragments*, the genome sequences situated between the high confidence genes. Genes predicted by GeneMark.hmm in the intermediate fragments are used as seeds to find homologous proteins in the database. These homologs are then mapped back to the genome with ProtHint (Brůna et al., 2021) to generate *hints* on the gene structure that are integrated into another round of the exon-intron structure prediction. GeneMark-ETP runs iterations of training, hint generation and gene prediction. It outputs the high confidence genes, further genes predicted by GeneMark.hmm in intermediate fragments and the hints from the proteins and RNA-seq.

The data for generating external hints is processed by GeneMark-ETP. Similar to GeneMark-EP+, intron hints, as well as start and stop codon hints, are created by ProtHint from spliced alignments of database proteins to the genome [Bruna et al., 2021]. Similarly to GeneMark-ET, intron hints are created by spliced alignments of RNA-seq to genome by HISAT2 [Kim et al., 2019]. Notably, a new type of external hints is created from assembled StringTie2 transcripts [Kovakaet al., 2019]. Protein-coding genes are predicted in assembled transcripts by GeneMarkS-T [Tang et al., 2015]. The level of confidence in such a prediction is determined on the basis of the alignment of the predicted protein to the proteins in the reference database. Out of these predictions, we select those that have high similarity scores. Besides these types of gene predictions, some other genes predicted in transcripts are selected based on the quality of *ab initio* predictions and other criteria as described in the description of GeneMark-ETP [Bruna et al., 2023b]. This set of high-quality gene predictions in assembled transcripts gives rise to a set of High Confidence (HC) genes. Overall, GeneMark-ETP creates three distinct groups of the external hints: external hints with both transcript and protein similarity support, hints with transcript and *ab initio* support and hints supported by protein similarity only (generated by ProtHint). All these sets are used for training of the statistical model and expanding of the set of HC genes to a set of genome wide gene predictions by GeneMark-ETP.

At the next step, AUGUSTUS is trained on the set of high confidence genes and predicts a second genome wide gene set with the support of the hints. At the final step, an updated TSEBRA (described next) combines the predictions made by AUGUSTUS and GeneMark-ETP while integrating the high confidence genes directly into the result to ensure their inclusion. The workflow is illustrated in Figure 1.

**Figure 1:**
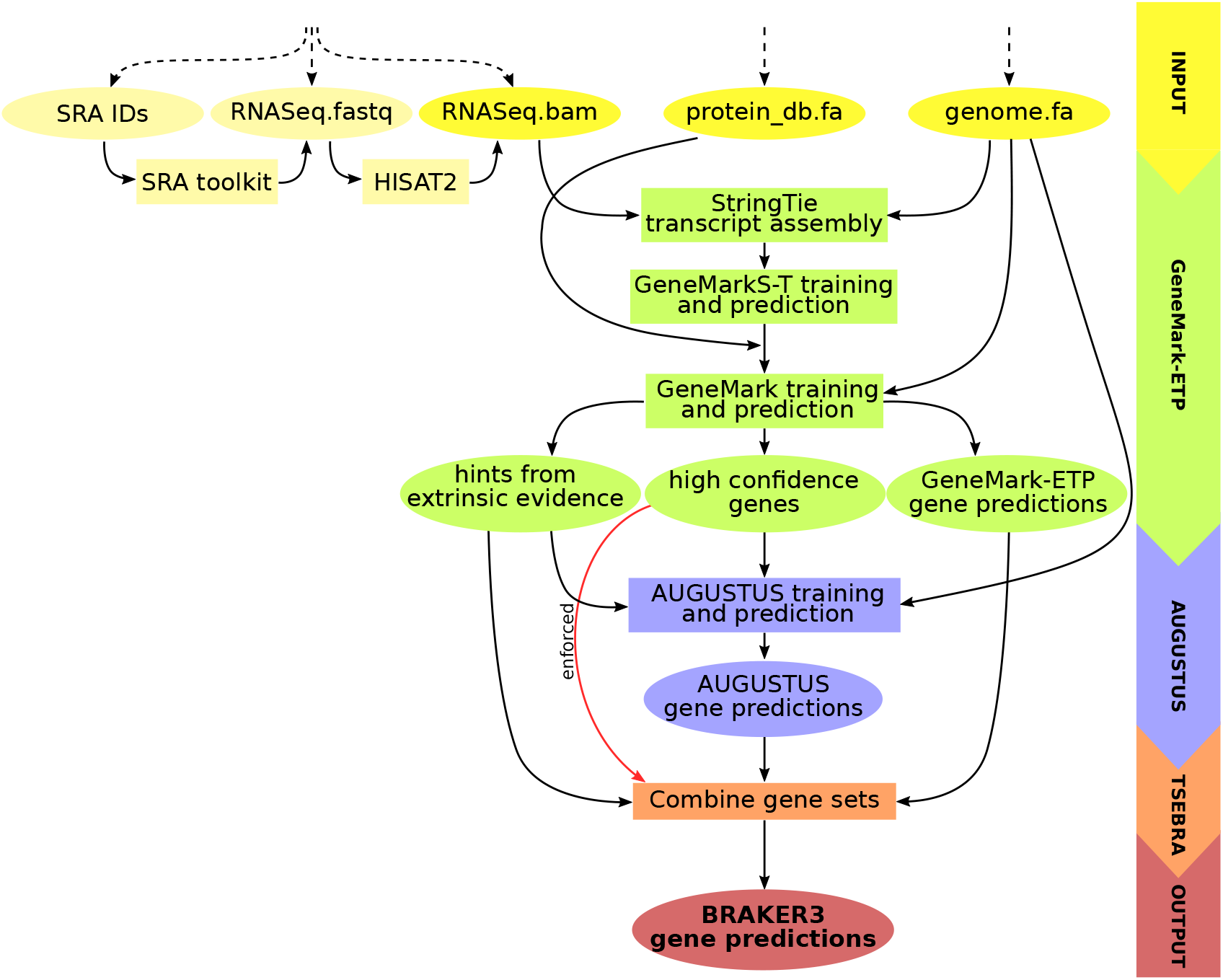
Schematic view of the BRAKER3 pipeline. Required inputs are genomic sequences, shortread RNA-seq data, and a protein database. The RNA-seq data can be provided in three different forms: IDs of libraries available at the Sequence Read Archive (Leinonen et al., 2010), unaligned reads or aligned reads. If library IDs are given, BRAKER3 downloads the raw RNA-seq reads using the SRA Toolkit (https://trace.ncbi.nlm.nih.gov/Traces/sra/sra.cgi?view=software) and aligns them to the genome using HISAT2 (Kim et al., 2019). It is also possible to use a combination of these formats when using more than one library.

### TSEBRA

The Transcript SElector for BRAker (TSEBRA) was improved and its original use in the BRAKER suite was extended. As described earlier, TSEBRA combines gene sets by evaluating and comparing candidate transcript isoforms using four transcript scores, which measured the agreement of transcript structures with extrinsic evidence. The extrinsic evidence is here utilized in the form of positions of supported exon borders, particularly intron position intervals. We have now introduced normalization of these transcript scores with respect to all input gene sets to TSEBRA, so the support with evidence is measured relative to the available evidence for the target genome. Normalization of a transcript score *s* for the *i*-th transcript of the input gene sets is defined as 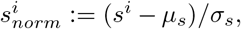 where *µ*_*s*_ and *σ*_*s*_ are the average and standard deviation of one of four transcript score measures *s*, calculated from scores of all transcripts in the input gene sets that TSEBRA is requested to combine. TSEBRA heavily relies on intron position information, which can make it challenging to evaluate single-exon transcripts. Therefore, the original TSEBRA tended to overestimate single-exon transcripts in some cases. To address this, we added a new option to TSEBRA that allows filtering out those single-exon genes that are predicted without any support by start- or stop-codon hints. When run by BRAKER3 on genomic sequences longer than 300Mbp, TSEBRA removes such single-exon genes that are predicted purely *ab initio*.

We also added TSEBRA to the workflow of BRAKER1 and BRAKER2, where it is now used to combine AUGUSTUS predictions with transcripts from GeneMark-ET/EP that are highly supported by extrinsic evidence.

### Test data

To benchmark BRAKER3 we selected 11 species: *Arabidopsis thaliana, Bombus terrestris, Caenorhabditis elegans, Danio rerio, Drosophila melanogaster, Gallus gallus, Medicago truncatula, Mus musculus, Parasteatoda tepidariorum, Populus trichocarpa* and *Solanum lycopersicum*. For each species, we retrieved: genome assemblies, 5 or 6 randomly selected short-read RNA-seq libraries from NCBI’s Sequence Read Archive (detailed list in Supplemental Table S15), a protein database, and a reference genome annotation (detailed list in Supplemental Table S1). Before running the experiments, we soft-masked repeats in the genomic sequences using RepeatModeler2 (Flynn et al., 2020). All data was publicly available. Genome versions, repeat masking and annotation processing are documented at https://github.com/gatech-genemark/EukSpecies-BRAKER2 and at https://github.com/gatech-genemark/GeneMark-ETP-exp.

For each target species, we prepared three differently sized protein databases, here termed *species excluded, order excluded* and *close relatives included*. The first two types of databases contain proteins from OrthoDB of species from the same broad taxonomic clade as the target, e.g. *Arthropoda* for *Drosophila melanogaster*. For this, OrthoDB was partitioned into the subsets of proteins for *Arthropoda, Metazoa, Vertebrata* or *Viridiplantae*. In the *species excluded* set of protein databases, we excluded for each target species all proteins from OrthoDB of that very species. In the *order excluded* databases, we removed for each target species all proteins of the same order as the target species. With these two large databases we test settings in which most of the possibly useful available proteins are used as informants. For the *close relatives included* set of databases, we selected for each species a small number of 4-12 closely related species and included their complete proteomes (Supplemental Table S2). These databases are much smaller than the corresponding *species excluded* and *order excluded* databases, by a factor between 17 and 132. The *close relatives included* databases were used to compare the BRAKER3 performance with performances of the other genome annotation tools that could not handle or were not designed to use larger databases: Funannotate failed to run on most of the large OrthoDB-based protein databases and MAKER2 was designed to be used with a smaller protein database, too.

It should be noted that the species for the *close relatives included* database were manually selected and the procedure would not scale well when very large numbers of species are annotated.

Above listed 11 benchmark species were sequenced relatively early in their respective clade and their annotations may have been used when annotating others. To avoid sampling bias associated with this choice, we also selected three recently sequenced genomes that do not have an established reference annotation: *Prunus dulcis* (almond), *Thrips palmi* (hemimetabolous insect), *Tetraselmis striata* (green algae).

### Experiments

We evaluated the performance of BRAKER3 and compared it with seven other methods: the previous versions BRAKER1 and BRAKER2 using only one type of extrinsic evidence (as included in the BRAKER v3.0.2 suite), TSEBRA (v1.1.0), combining the results of BRAKER1 and BRAKER2, MAKER2 (3.01.04), FINDER (v1.1.0), Funannotate (v1.8.14), and GeneMark-ETP. As BRAKER3, BRAKER2, FINDER and GeneMark-ETP can use a large protein database and since doing so saves a manual step we compared these four tools along with BRAKER1 and TSEBRA in two sets of experiments, where the large *order excluded* and *species excluded* databases were used. In another set of experiments, we compared BRAKER3 with MAKER2 and Funannotate on the smaller and target-specific *close relatives included* databases using the same RNA-seq data as in the other experiments.

When running Funannotate, we tried two recommended flags for generating gene sets, a specific handling of repetitive regions and an additional gene model update step. This resulted in four variant sets of gene predictions per genome. Here, we report the numbers of the variant of Funannotate that performed best (both flags were set, Supplemental Table S11).

MAKER2 was executed according to recommendations provided by the developers of MAKER2, integrating GeneMark, AUGUSTUS and SNAP predictions. The details are provided in Supplemental Table S16. MAKER2 does not provide automatic training procedures. A recommended approach is the manual execution of training runs of all the *ab initio* programs outside of MAKER2. To provide the best possible models, we trained SNAP and AUGUSTUS on the respective reference annotation which all programs were evaluated on, unless models for SNAP or AUGUSTUS for the species were included in the standard distribution of these tools. Models for GeneMark were also chosen to match the best possible training routine (see Supplementary Material). This approach allowed for the automatic execution of MAKER2. However, the quality of the trained parameters of the gene finders we used for MAKER2 can be considered rather as *upper limits of what can be expected* on new genomes.

We compared the predicted genome annotations with the reference annotations to assess the performance of BRAKER3 on exon, gene and transcript levels. This long-established approach to benchmark gene prediction methods allows to measure grave as well as subtle errors and is informative in absolute terms if the reference annotations are of high quality. Note that, particularly for model species such as *Mus musculus* and *Drosophila melanogaster*, diverse sources of evidence besides RNA-seq have contributed to the manually supervised annotation of these genomes. This includes outcomes from functional studies, such as knockdown experiments, and the findings of comparisons between multiple genomes (Lilue et al., 2018). Further, the available evidence is vast for some model species, e.g. the Sequence Read Archive includes more than a million RNA-seq libraries for *Mus musculus* and more than 80k RNA-seq libraries for *Drosophila melanogaster*. Therefore, to estimate the performance on new genomes which typically have much less available evidence, we deliberately limited the evidence and used RNA-seq only from at most 6 libraries and no closely related proteins. It should be noted that some of the reference annotations may fall short of the desired accuracy, potentially leading to an underestimation of accuracy or a bias in our analysis.

As metrics, we used the *sensitivity* (Sn=TP/(TP+FN)) - the percentage of correctly found instances from the reference annotation, the precision (Prec=TP/(TP+FP)) - the percentage of correct instances in the predicted annotation, and the F1-score - the harmonic mean of Sn and Prec.

Note that most previous publications on gene finding methods and their benchmarking have used the term ‘specificity’ to refer to the accuracy measure defined as TP/(TP+FP). However, in the related and fast-growing field of machine learning, the term ‘specificity’ has the meaning of true negative rate TN/(TN+TP). To unify the designations, we switch to the use of the term ‘precision’ for TP/(TP+FP) and utilize this designation in all the figures and tables in this manuscript.

When evaluating on exon level or transcript level, each transcript / exon was individually assessed. However, when evaluating on gene level, a predicted gene was counted as true positive if at least one of its predicted alternative transcripts matched a reference transcript.

To assess BRAKER3 on the three recently sequenced genomes without using a reference annotation we used BUSCO Simão et al. (2015). BUSCO was run to assess the ‘completeness’ of genome annotations performed with BRAKER3. The BUSCO completeness score is an estimate of the percentage of genes, that are generally present in the respective clade in a one copy, that are found. BUSCO version 5 was run with the database version 10.

## Results

### Assessment of performance of BRAKER3

For each species, computational experiments were done by running five gene prediction tools, BRAKER1, BRAKER2, TSEBRA, GeneMark-ETP and BRAKER3. These tools were run on each genome with extrinsic information in the form of species specific set of RNA-seq libraries and two types of species specific protein databases, the order excluded and the species excluded (see Test data section). The quality of the annotation depends generally on the evolutionary relationship of the species whose genome a user may want to annotate (the *target*) to those species that have well-established genome annotations. To give a range of performance estimates, we performed experiments with the particularly favorable case of the *species excluded* database and with the rather conservative assumption of the *order excluded* database. We show the averaged accuracy measures (Sn and Prec) at exon, gene and transcript level of BRAKER3 and four other gene finding tools on the 11 genomes, with species-specific *order excluded* databases (Figure 2). The pipelines in order of increased performance are BRAKER1, BRAKER2, TSEBRA combining BRAKER1 and BRAKER2, GeneMark-ETP and BRAKER3. Detailed information for each genome is given in Supplemental Tables S4 and S5. A species-by-pipeline heatmap of F1-scores at gene level is shown in Supplemental Figure S1.

**Figure 2:**
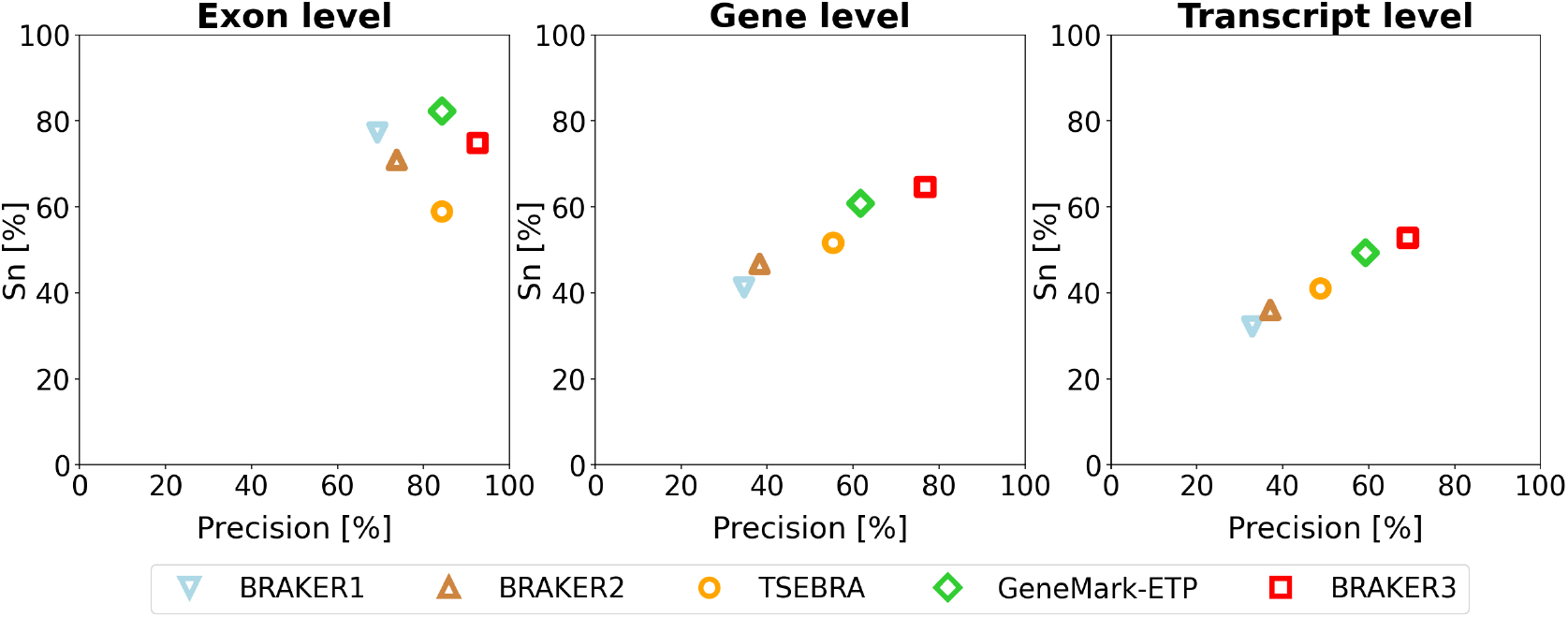
Average precision and sensitivity of gene predictions made by BRAKER1, BRAKER2, TSE-BRA, GeneMark-ETP, and BRAKER3 for the genomes of 11 different species (listed in Supplemental Table S1). Inputs were the genomic sequences, short-read RNA-seq libraries and protein databases (*order excluded*).

Notably, there was a significant improvement of BRAKER3 in comparison with BRAKER1 and BRAKER2 in species with GC-heterogeneous or large genomes (Figure 3). The highest performance increase was achieved in *Gallus gallus*, where the BRAKER3 F1-score on gene / transcript level was improved by 55/48 points compared to the combined prediction of BRAKER1 and BRAKER2 generated by TSEBRA (Supplemental Table STable 4).

**Figure 3:**
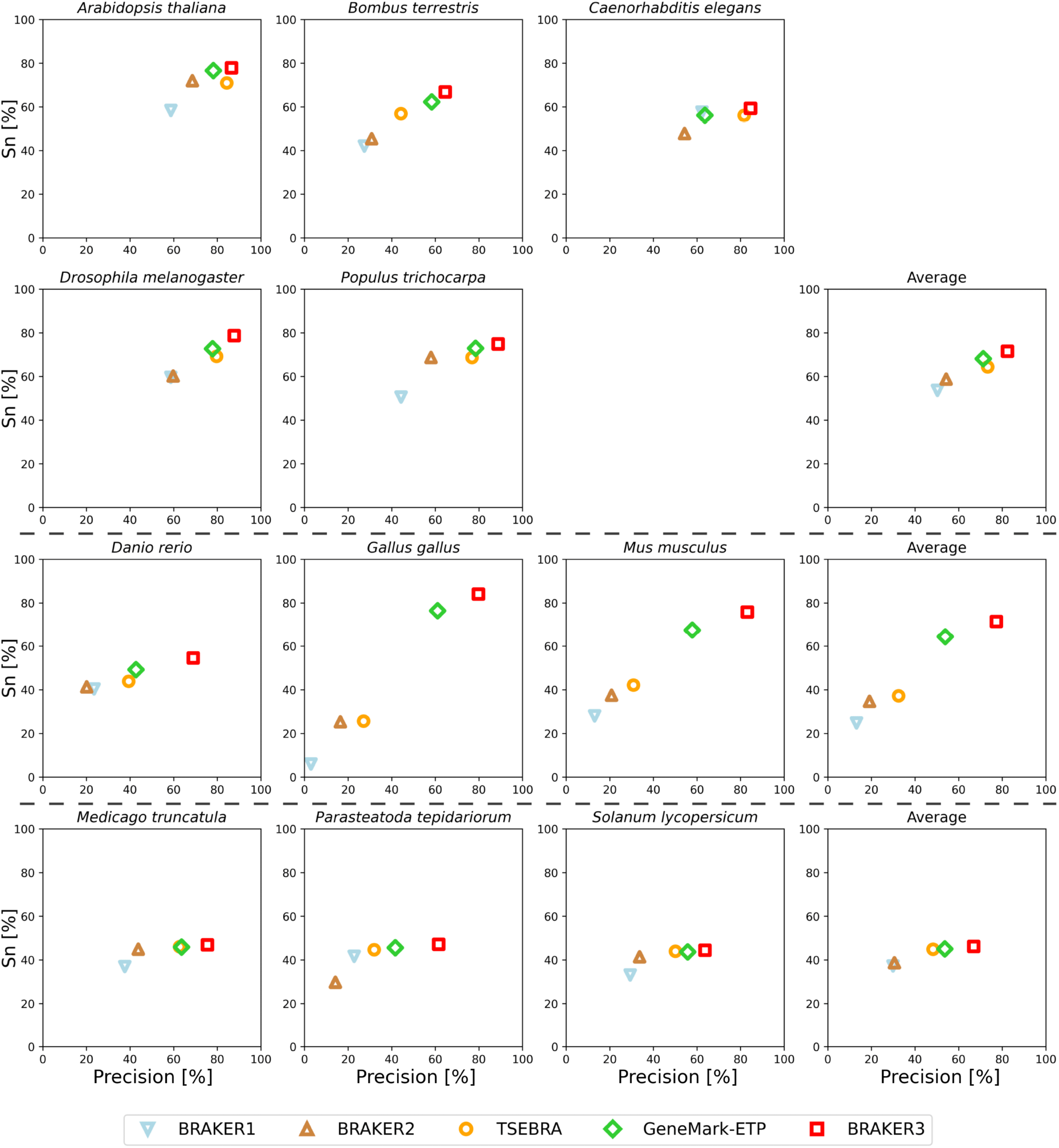
Gene level precision and sensitivity of gene predictions made by BRAKER1, BRAKER2, TSEBRA, GeneMark-ETP, and BRAKER3 for the genomes of 11 different species: well annotated and compact genomes (first and second row), well-annotated and large genomes (third row), other genomes (fourth row). The fourth column shows the average for each group. Inputs were the genomic sequences, short-read RNA-seq libraries, and protein databases (*order excluded*).

Here, BRAKER3 greatly benefited from the high accuracy of GeneMark-ETP and managed to exceed the sensitivity and precision on gene and transcript levels even further. GeneMark-ETP enabled the generation of a highly precise set of high confidence genes to train the AUGUSTUS model. As a result, this AUGUSTUS prediction using extrinsic evidence of BRAKER3 had a higher sensitivity than GeneMark-ETP on gene, transcript and exon level at the cost of lower precisions. AUGUSTUS’ average gene and transcript level F1-scores of 59.6 and 51.3, respectively, exceeded the F1-scores of the AUGUSTUS predictions in BRAKER1 and BRAKER2, and are slightly lower than the F1-scores of GeneMark-ETP, see Supplemental Table S10. By integrating TSEBRA into BRAKER3 and combining sets of gene predictions made by GeneMark-ETP and AUGUSTUS, the final BRAKER3 predictions achieved higher sensitivity and precision than either GeneMark-ETP and AUGUSTUS at both the gene and transcript level. Further, the BRAKER3 annotation has higher precision on exon level than the annotation of GeneMark-ETP and the AUGUSTUS annotation that BRAKER3 produces for all the 11 species (Supplemental Tables S4 and S10). TSEBRA tends to eliminate false transcripts from either input annotation, one such example is shown in Supplemental Figure S2.

BRAKER3 is more likely to make an error when predicting an unspliced coding region, i.e. a gene that has a single coding sequence (CDS) feature, than when predicting a CDS of a multi-CDS gene (Supplemental Figure S3). It is even the case that for 9 out of 11 species the transcript-level F1-score for predicting *spliced* transcripts is larger than the respective score for unspliced transcripts (Supplemental Figure S4). This may come as a surprise as the potential for predicting differences in two gene structures is larger for multi-exon genes and with any difference the predicted transcript is counted as false. These findings agree with previous research indicating that gene finders generally show decreased performance on unspliced transcripts (Scalzitti et al., 2020). This reduced performance could be attributed to the inherent design of the models or a lack of representation of single-exon genes in the training datasets. In spliced genes, BRAKER3 has more difficulties to predict the initial CDS that contains the start codon than the terminal CDS that contains the stop codon. The highest exon level F1 score of about 88% is achieved for internal CDS (Supplemental Figure S3). BRAKER3 predicts acceptor and donor splice sites equally well with an F1-score of *>* 87% averaged over all species. The averaged F1-score for stop and start codons are 76% and 70%, respectively (Supplemental Figure S5).

In all species, transcripts are much more likely to be correctly identified by BRAKER3 if they are supported by more RNA-seq reads. Figure 4 shows the transcript level sensitivity for three terciles of expression levels, measured using the RNA-seq libraries that were used for prediction as well. When averaging over all species, only 23% of low-expression transcripts are correctly identified, 55% of medium-expression transcripts, and 76% of highly expressed transcripts. Note that there are multiple explanations or factors that may contribute to this observation. BRAKER3 and reference annotations may be more accurate for transcripts that have more RNA-seq support, directly as a consequence of this evidence. However, highly expressed genes may also be better represented by statistical models of gene finding, e.g. because preferred codons may make translation more efficient (Hershberg and Petrov, 2008).

**Figure 4:**
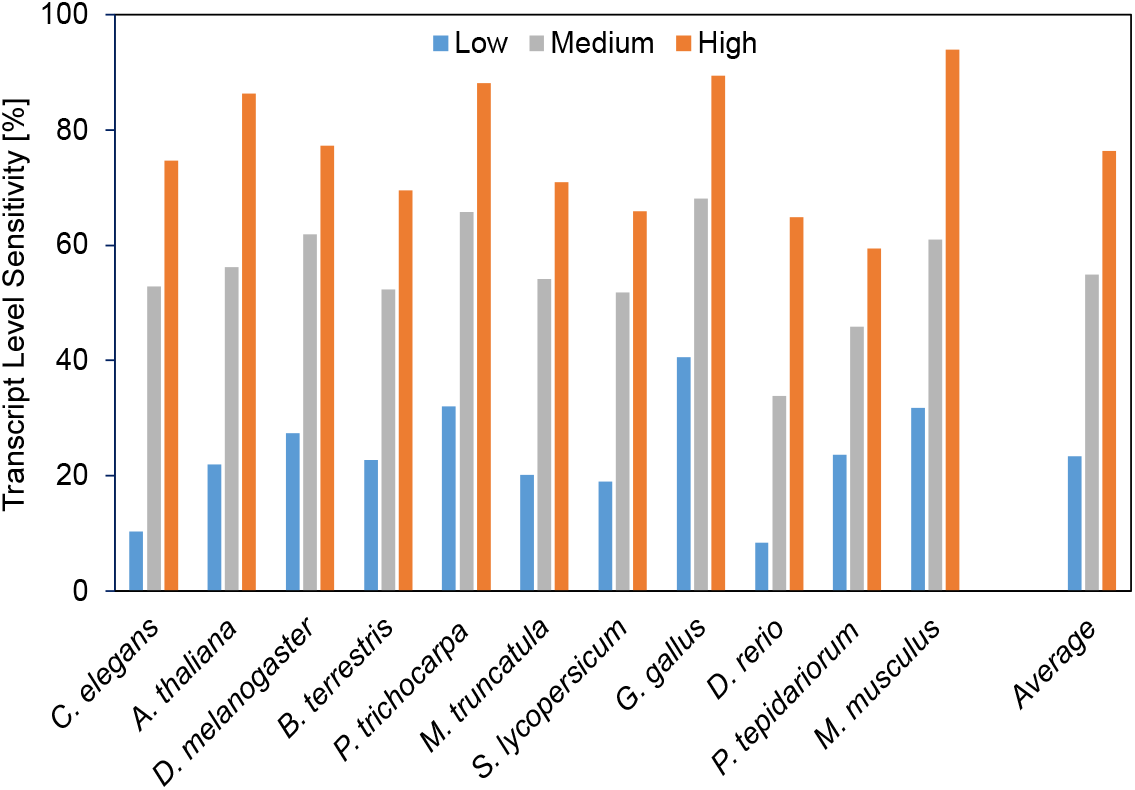
Low, medium and highly expressed transcripts are in the first, second and third tercile of expression levels, respectively.

When we used the *species excluded* protein database, which may include very closely related species, the performance measures of the methods using the protein data increased overall (Supplemental Table 6). On average, the BRAKER3 transcript level sensitivity was improved by approximately 3 percentage points and the precision was improved slightly (less than 1 percentage point). However, the relative ranking of the methods and the comparison of BRAKER3 with other methods remain unchanged.

BRAKER3 had the highest sensitivities and precisions for each species at the transcript and gene level, but often had a somewhat lower exon-level F1-score than GeneMark-ETP (Supplemental Table S5). In each species, BRAKER3 was more precise in predicting exons than GeneMark-ETP, which in turn predicted exons more sensitively than BRAKER3 (Supplemental Table S4). Thus, there was a trade-off in exon sensitivity and precision between the two methods, with an average difference of approximately 8 percentage points in both measures (Supplemental Table S4). We presume that the occasional false-positive exons of GeneMark-ETP hurt the stricter transcript and gene performance measures more than those exons occasionally missed by BRAKER3 do.

The set of transcripts found by BRAKER3 and GeneMark-ETP, respectively, have large overlaps (Supplemental Figure S6). Transcripts uniquely predicted by BRAKER3 and not by GeneMark-ETP uncover more of the remaining reference annotation transcripts than vice versa. This pattern is consistent across all 11 species and applies to both spliced and unspliced transcripts. TSEBRA selects most of the single-exon genes predicted by GeneMark-ETP to be in the final set of BRAKER3 genes (Supplemental Figure S6). However, it adds single-exon genes predicted by AUGUSTUS, which increases the percentage of single-exon genes correctly identified by 5.5 percent points on average. In the particular case of *C. elegans*, even about half of the single-exon genes predicted correctly by BRAKER3 are predicted by AUGUSTUS alone (Supplemental Figure S6).

Supplemental Figure S7 breaks down the sensitivity with which transcripts are correctly identified by expression level and quantifies how AUGUSTUS and GeneMark-ETP complement each other when run in BRAKER3. The BRAKER3 transcript sensitivity benefits the most from the integration of AUGUSTUS for medium-expressed transcripts. In this expression tercile, on average 8.2% of the transcripts are identified by BRAKER3 (and AUGUSTUS) but not by GeneMark-ETP. Remarkably, of the medium-expressed mouse transcripts BRAKER3 correctly identifies about 16% more than GeneMark-ETP. A possible explanation of this observation is that highly expressed transcripts can be identified well by GeneMark-ETP, e.g. by searching them in assembled transcripts, and both GeneMark-ETP and AUGUSTUS still have trouble correctly predicting low expressed transcripts (Supplemental Figure S7).

### Comparison of BRAKER3 to MAKER2, Funannoate and FINDER

BRAKER3 was compared with MAKER2 and Funannotate on eight of the 11 genomes used in the previously described tests. The relatively large genomes of *Mus musculus* and *Parasteatoda tepidariorum* (errors in the Funannotate runs) and *Danio rerio* (error in the MAKER2 run) were excluded because Funannotate or MAKER2 failed to finish even for the smaller *close relatives included* protein sets.

Figure 5 and Supplemental Table S8 show the comparison of BRAKER3 to MAKER2 and Funannotate. All tools, including BRAKER3, are given as input the smaller *close relatives included* protein databases and the same RNA-seq data as in all experiments. BRAKER3 consistently outperforms Funannotate and MAKER2 on exon, gene and transcript level (Figure 5). On average, BRAKER3’s F1-scores were higher than the ones of Funannotate by 10.2 points at the exon level, 25.9 points at the gene level and 21.6 points at the transcript level. In turn, Funannotate exceeded MAKER2 by 2.2, 3.8 and 4.4 points with regards to the F1 measure on exon, gene and transcript level, respectively. BRAKER3 shows better performance than Funannotate and MAKER2 for all species and individual metrics, except at the exon level for *Caenorhabditis elegans*, where BRAKER3 had a sensitivity 3.3 percentage points lower than the one of Funannotate (Supplemental Table S8).

**Figure 5:**
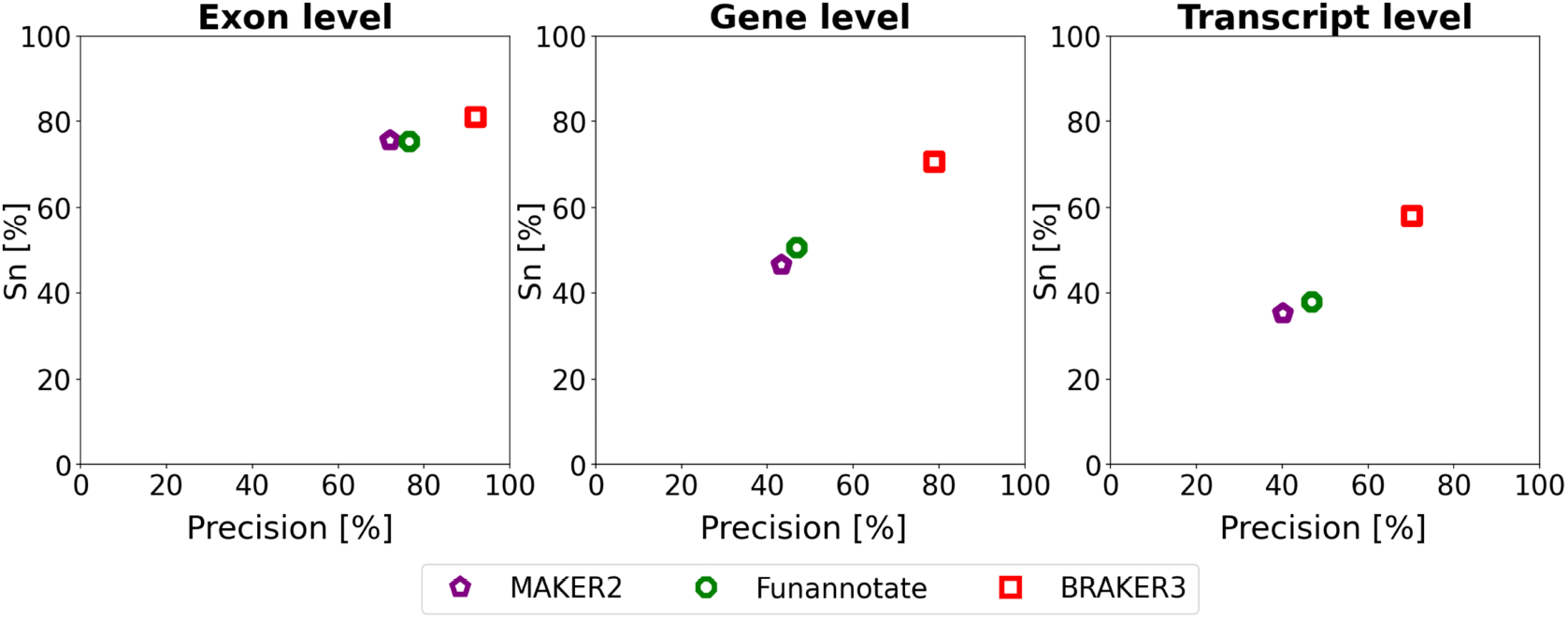
Average precision and sensitivity of gene predictions made by MAKER2, Funannotate, and BRAKER3 for a subset of 8 species (excluding the mouse, spider and fish genome). Inputs were the genomic sequences, short-read RNA-seq libraries, and protein databases (*close relatives included*). The accuracy of MAKER2 reported here can be regarded as an upper limit of what can be expected when annotating a previously unannotated genome (see Experiments section).

We compared the results of BRAKER3 on the protein informant databases *close relatives included* and *species excluded*. Both series of databases may contain proteins from close relatives of the target, but the database *close relatives included* is much smaller. When run with the *species excluded* database (Supplemental Tables S6 and S7) BRAKER3 has on average an F1-score that is higher by 0.40, 0.23 and 0.34 on the exon, gene and transcript level, respectively, than when BRAKER3 is run with the *close relatives included* database (Supplemental Tables 8). Thus, when BRAKER3 uses the larger protein database it delivers slightly better results. Perhaps more importantly, using the broader database has a practical advantage in that it does not require the (manual) step to compile a database of closely related proteomes.

The FINDER annotation pipeline was run on the *order excluded* databases with the same input data as BRAKER3, but the execution only completed for 7 of 11 species. However, its best performance, a *∼*15 gene F1-score and a *∼*11 transcript F1-score for *Drosophila melanogaster*, was much below the respective values of the other methods (Supplemental Tables S9). A reason that the performance of FINDER was below the figures published by Banerjee et al. (2021) could be that Banerjee et al. did not report the exclusion of any proteins from the UniProt informant database. Also, Banerjee et al. used a much larger set of RNA-seq libraries.

### BUSCO gene set completeness

We ran BRAKER3 also on the three recently sequenced genomes *Prunus dulcis, Thrips palmi* and *Tetraselmis striata*. BUSCO was used to estimate the completeness of the 11 + 3 = 14 predicted proteomes (databases listed in Supplemental Table S3). The average BUSCO completenesses (single copy or duplicate) of BRAKER3 are 96% for the group of well-annotated and compact genomes, 83.6% for the group of well-annotated and large genomes, 91% for the group of other genomes with a reference annotation (*M. truncatula, P. tepidariorum, S. lycopersicum*) and 93.7% for the group of 3 recent genomes (Supplemental Figure S9). The figure also shows the BUSCO scores of AUGUSTUS and GeneMark-ETP and the respective total numbers of genes. For each of the groups of species, AUGUSTUS had a higher BUSCO completeness score than GeneMark-ETP, which in turn had a higher score than BRAKER3. However, this is also the order of tools sorted by decreasing numbers of predicted genes, i.e. there is a – perhaps unsurprising – tendency that a tool that predicts more genes genome-wide also reports more of the universal single copy genes that are counted by BUSCO. When comparing the gene precisions, here the percentages of predicted genes that exactly share at least one transcript with the reference annotation, we observe the reversed order of tools: BRAKER3 has a lower percentage of false positive genes than GeneMark-ETP which has in turn a lower percentage than AUGUSTUS (Supplemental Figure S10). The quantitative view of this issue varies between species and groups. As a trend, a relatively large advantage of BRAKER3 in gene precision stands against a relatively small advantage of AUGUS-TUS in BUSCO completeness. For example, in the group of well-annotated large genomes, AUGUSTUS is 6.6 percent points more BUSCO complete than BRAKER3, but BRAKER3 is 30.5 percent points more precise than AUGUSTUS. In conclusion, for these gene sets, a better BUSCO score comes at the expense of more false positive genes, and BRAKER3 weights the trade-off between fewer missing genes and fewer false positive genes in favor of the latter. We make a point that the BUSCO scores should be viewed as a rough measure of gene prediction sensitivity (see Discussion). Moreover, BUSCO uses a set of genes that are well-conserved and, therefore, arguably relatively easy to predict.

### Runtime

We ran all methods except MAKER2 on an HPC node with Intel(R) Xeon(R) CPU E5-2650 v4 @ 2.20GHz using 48 threads. The pipelines in order of runtime when using the *order excluded* databases were BRAKER1, GeneMark-ETP, BRAKER2, BRAKER3. The BRAKER3 runtime ranges from 5h 37min in *Arabidopsis thaliana* to 64h 16min in *Mus musculus*. The time for aligning the RNA-seq reads is not included in these figures. However, parameter training is an integral part of the pipeline and its duration is included. Despite having the longest run-time of all methods, BRAKER3 can annotate even large genomes in a reasonable time. The *order excluded* protein databases are roughly 1-2 orders of magnitude larger than the *close relatives included* protein databases (see Supplemental Table S1). Nevertheless, BRAKER3 required only 23% more time on these large protein databases (averaged over 8 species, compare Supplemental Tables S12 and S14). Figure 6 shows the runtime as a function of genome size and protein database choice. A linear regression of 19 BRAKER3 whole-genome runtimes yielded the estimate

**Figure 6:**
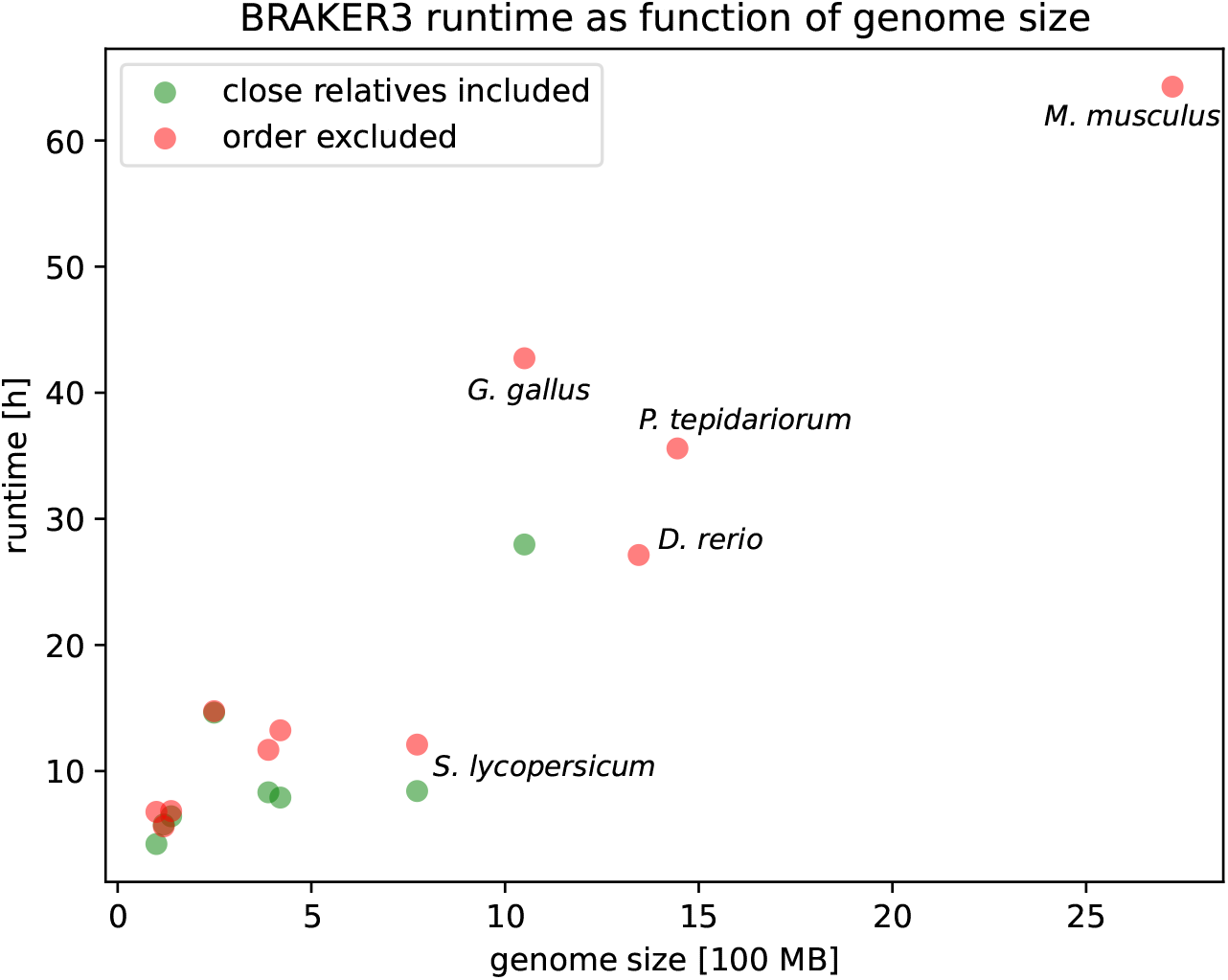
The execution time of BRAKER3. The time required for aligning the RNA-seq to the genome and thus producing the `BAM` input files is not included.

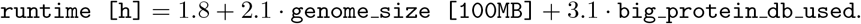

Here, `big protein db used` is 1 if the OrthoDB partition is used (*order excluded*) and 0 if the small *close relatives included* protein database is used. Consequently, using the large protein database adds an estimated 3.1 hours to the runtime. It should be noted that many factors influence runtime and the linear regression ansatz can only give a rough estimate. Supplemental Figure S8 shows a comparison of predicted to actual runtimes.

We also compared the runtimes of BRAKER3, Funannotate and MAKER2 on the smaller *close relatives included* protein databases. Funannotate required on average roughly the same time as BRAKER3 (see Supp. Table 14). As we ran MAKER2 on faster hardware (see Supplemental Material) we made a runtime comparison experiment with BRAKER3 on the same hardware for *Drosophila melanogaster*. When given the relatively small protein database as input (116,493 proteins) MAKER2 took 2.1 hours and BRAKER3 took 3.5 hours. When given a large protein database as input (2,588,444 proteins), MAKER2 took 16 hours and BRAKER3 took 2.5 hours. The run-time of BRAKER3 is much less dependent on the protein database size (here its decrease was due to a variable-duration hyperparameter optimization step during training). When comparing these runtimes, one has to consider that the figures for MAKER2 do not include the considerable times for training gene finders and neither for transcriptome assembly. In contrast, BRAKER3 performs these steps as part of the pipeline. Some further examples of runtimes of MAKER2 are shown in Supplemental Table S13.

### Virtualization

One problem with modern genome annotation pipelines is their dependence on an increasing number of tools, which can make their installation and maintenance difficult. Therefore, we provide a Docker container for BRAKER, making it easy to install and use.

## Discussion

We present BRAKER3, a novel genome annotation pipeline for eukaryotic genomes that integrates evidence from transcript reads, homologous proteins and the genome itself. We report significantly improved performance for 11 test species. BRAKER3 outperforms its predecessors BRAKER1 and BRAKER2 by a large margin, as well as publicly available pipelines, such as MAKER2, FINDER and Funannotate. The most substantial improvements are observed in species with large and complex genomes. Additionally, BRAKER3 adds a Docker container that also easily executes with Singularity (Kurtzer et al., 2017) to the BRAKER suite, which makes it more user-friendly and easier to install.

BRAKER3’s final and integrative step, TSEBRA, selects transcripts from sets of transcripts from multiple sources. One source is gene prediction in transcripts assembled from RNA-seq, and two other sources are the two HMM models that predict genes in the genome using different approaches to evidence integration and, in the case of AUGUSTUS, produce alternative transcript variants. BRAKER3 combines these sets of transcripts and can be seen as an ensemble learning approach that improves each of its inputs. In fact, the combined BRAKER3 transcripts have a higher F1-score than either of the combined GeneMark-ETP and AUGUSTUS transcript sets.

BRAKER3’s performance drops for transcripts that are weakly represented in the RNA-seq data. Notably, BRAKER3 cannot be used without RNA-seq evidence. When only protein evidence is available, we recommend to use BRAKER2 for small and medium sized genomes and GALBA (Brůna et al., 2023a) for large vertebrate genomes.

Currently, BRAKER3 predicts only protein-coding genes. Consequently, it makes a decision for each assembled transcript if it encodes a protein or not. Therefore, a future release of BRAKER may give users the option to output transcripts that were not classified as coding as candidates for lncRNAs.

With our diverse species selection it was not feasible to use RNA-seq sets that are comparable across different species. However, Stanke et al. (2019) investigated the impact of an RNA-seq sampling method on the performance of BRAKER1. In their study, a manual selection of libraries from different experiments and tissues was compared with an automatic sampling from diverse libraries for complementary data. The latter yields on average slightly better performances in BRAKER1 when the number of reads was similar. However, such conclusions do not necessarily carry over to BRAKER3. This is because the assembly of the transcriptome used in BRAKER3 could benefit from more homogeneous data.

In our study, we selected well-annotated genomes as benchmarks to predict gene structures, inadvertently introducing a sampling bias, arguably towards species with relatively many related sequenced genomes. We corrected for this bias by restricting the amount and usefulness of available evidence. To address the issue of overrepresented annotated species closely related to our benchmark species, we eliminated homology evidence from the same taxonomic order. For instance, in annotating *D. melanogaster*, we used *Lepidoptera* species, such as butterflies, as the nearest relatives, providing a conservative approximation of the nearest annotated genomes for a typical new target genome. Furthermore, despite the availability of numerous RNA-seq libraries (for example, over 80,000 for *D. melanogaster*), we limited the input for BRAKER3 to a maximum of six RNA-seq libraries. To control for any remaining effects of the sampling choice on the relative assessment of the tools, ideally, one would sample test genomes that represent the *whole space of eukaryotic genomes*. However, a comprehensive gold standard annotation for such a set of genomes does not exist.

Although BUSCO is commonly employed as a standard for assessing the completeness of a proteome, we argue that it does not provide a comprehensive comparison with a mature reference annotation. Specifically, BUSCO’s design does not encompass the detection of various gene structure errors, nor does it impose penalties for the presence of false positive genes. Therefore, by itself BUSCO cannot be used to fully evaluate the accuracy and integrity of gene structures.

As long as the error rates in eukaryotic gene annotation remain significant, a trade-off between the numbers of false positive and false negative genes has to be made. The implications and severity of either type of error depend on the type of analysis conducted with the gene set. In comparative studies, for instance, false negative genes can lead to false hypotheses about missing genes, while false positive genes can lead to false hypotheses about orphan or taxonomically unique genes. Moreover, such false positives have the potential to erroneously influence annotations in related species and can at worst lead to failed experiments based on the assumed existence of these genes. In contrast to prioritizing sensitivity or achieving high BUSCO completeness scores, BRAKER3 focuses on the precision of its gene predictions. However, for users seeking higher sensitivity, the gene sets produced by GeneMark-ETP and AUGUSTUS might be more suitable choices.

## Software Availability

BRAKER3 is available on GitHub (https://github.com/Gaius-Augustus/BRAKER), as a Docker container (https://hub.docker.com/r/teambraker/braker3) and a Supplemental Code file. BRAKER3 and AUGUSTUS are distributed under the Artistic License. GeneMark-ETP and its part GeneMark.hmm, are distributed under the Creative Commons Attribution NonCommercial ShareAlike licence.

## Acknowledgements

The research was supported in part by the US National Institutes of Health grant GM128145 to MB and MS. The funding body did not play any role in the design of the study, in the collection, analysis, or interpretation of data or in writing the manuscript. AL and MB acknowledge a generous support of Microsoft who provided Azure cloud computing credits via Georgia Tech Cloud Hub.

## Author contributions

L.G., T.B., K.J.H., A.L., M.B., and M.S. conceptualized the pipeline, L.G. implemented BRAKER3, K.J.H. created the containers, L.G. and A.L. performed BRAKER and BUSCO experiments, A.L. and M.E. ran MAKER2 and FINDER, respectively, L.G., M.S. and A.L. created the figures, L.G, M.S. and M.B. wrote or edited the paper, with input from all authors.

