## Supplemental Material for "BRAKER3: Fully automated genome annotation using RNA-seq and protein evidence with GeneMark-ETP, AUGUSTUS and TSEBRA"

### 1 Supplemental Material

#### 1.1 Supplemental Figures

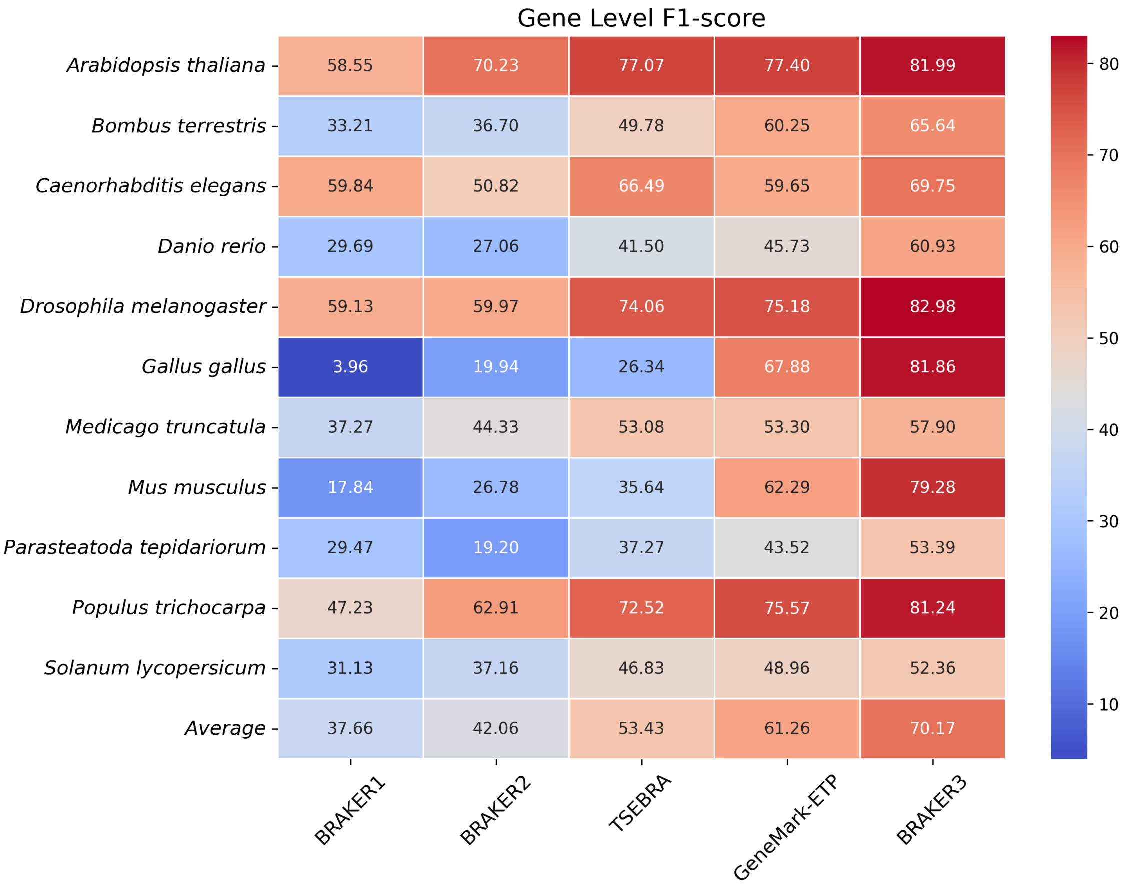


Supplemental Figure S1: Heatmap of F1-scores of pipelines being input short-read RNA-seq libraries and a protein database (with proteins of species from the respective **order excluded**). The last row shows the averages for the 11 different species.

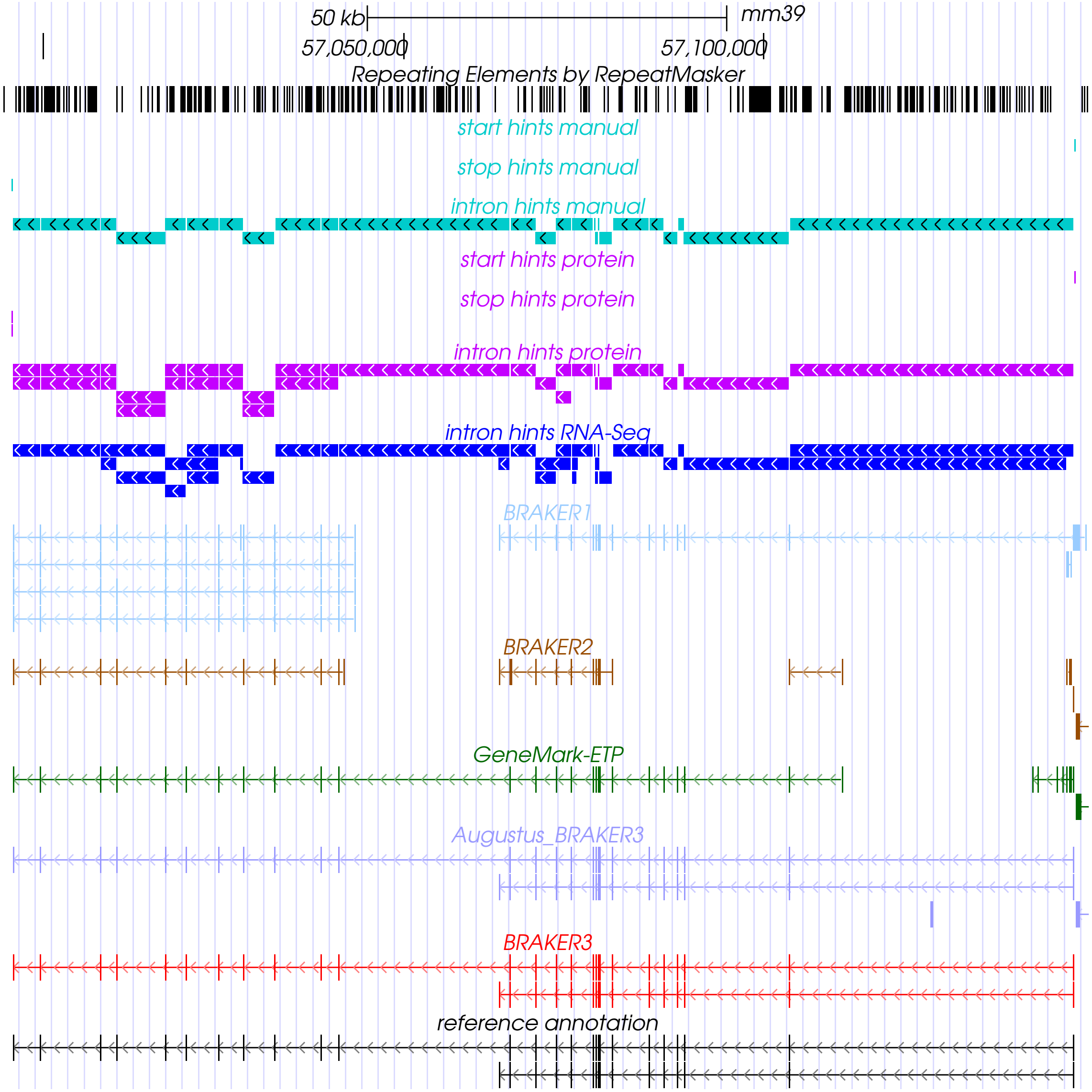


Supplemental Figure S2: A gene locus of *Mus musculus* visualized with the UCSC Genome Browser (Kent et al. (2002), `http://genome.ucsc.edu`). Tracks for the gene prediction of pipelines obtaining short-read RNA-seq libraries and a protein database (respective *order excluded*) as input are displayed. Tracks highlighting exon boundaries based on extrinsic evidence are shown at the top. Manual hints represent high confidence hints that are highly weighted during AUGUSTUS prediction. In this example, TSEBRA selects two correct transcripts from AUGUSTUS run as part of BRAKER3 (AUGUSTUS_BRAKER3) and filters out three false transcripts from GeneMark-ETP and two from AUGUSTUS.

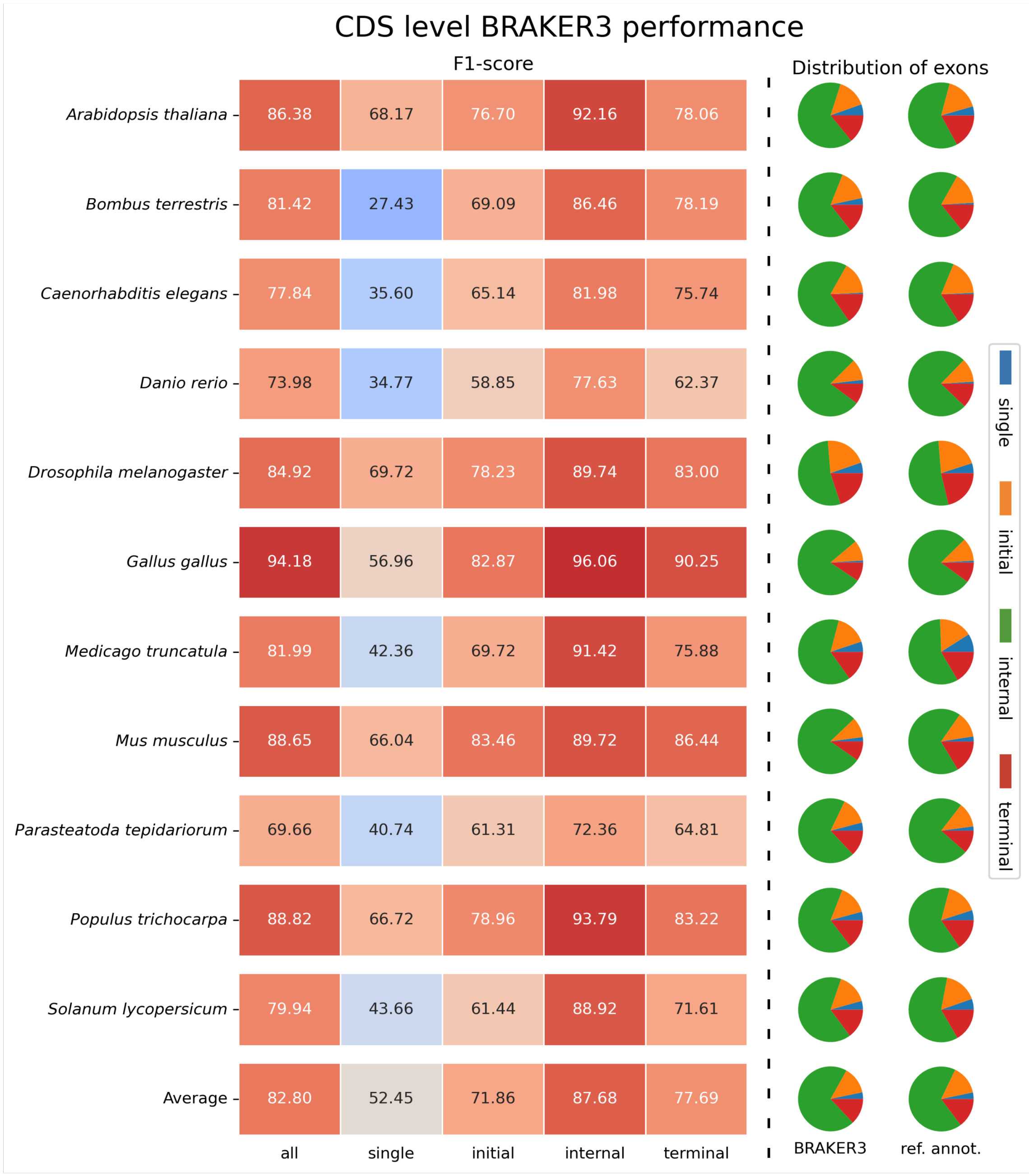


Supplemental Figure S3: The F1 score when predicting the protein-coding (parts of) exons, broken down by type of exon. Single: unspliced coding region, initial/terminal: first/last coding region, respectively, of a gene with spliced coding sequence, internal: coding region bordered by a splice site on both sides. The pie charts visualize the proportions of exon types in the BRAKER3 and reference annotations.

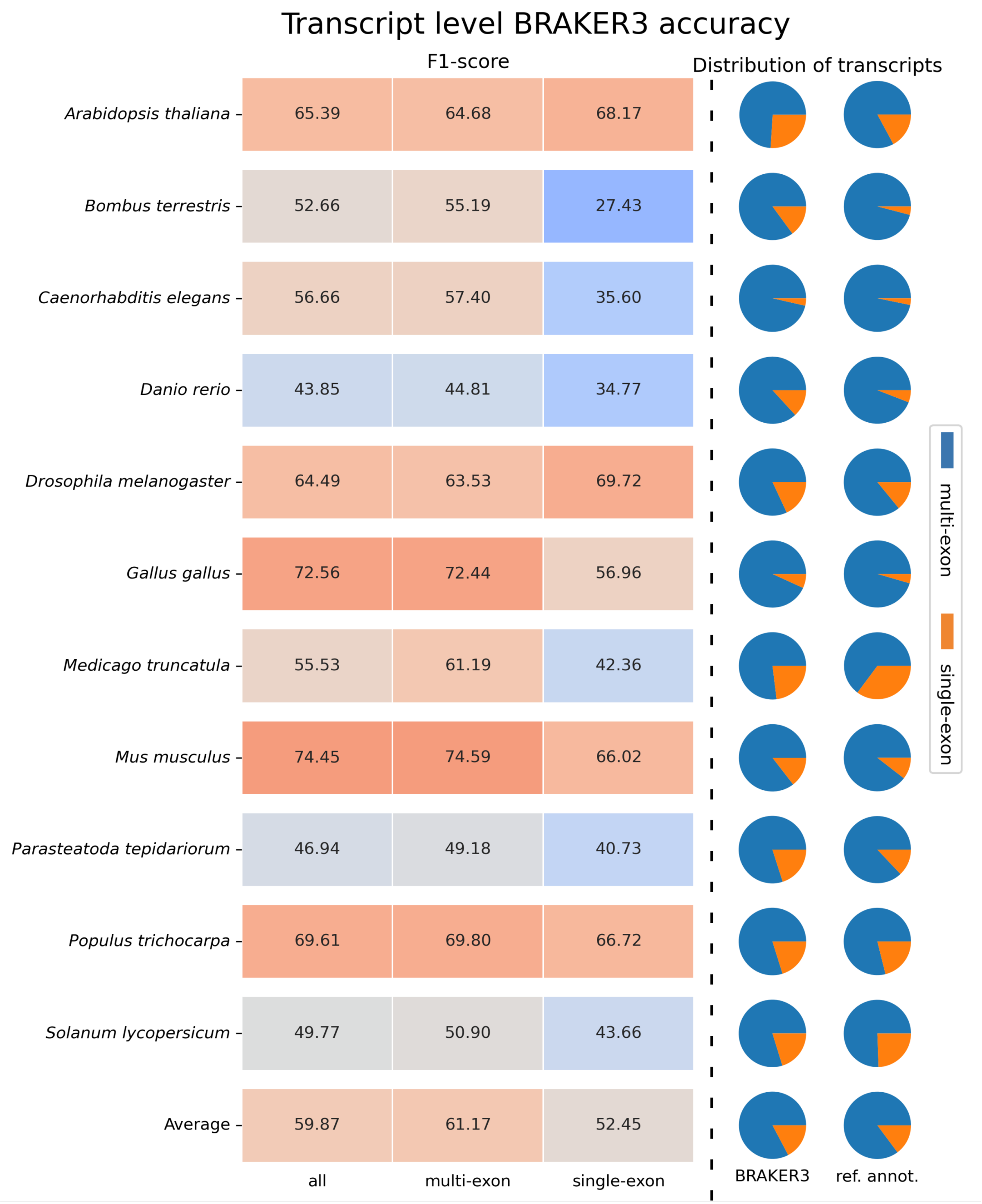


Supplemental Figure S4: Performance of BRAKER3 broken down by whether the coding sequence of a transcript is spliced (multi-exon) or unspliced (single-exon).

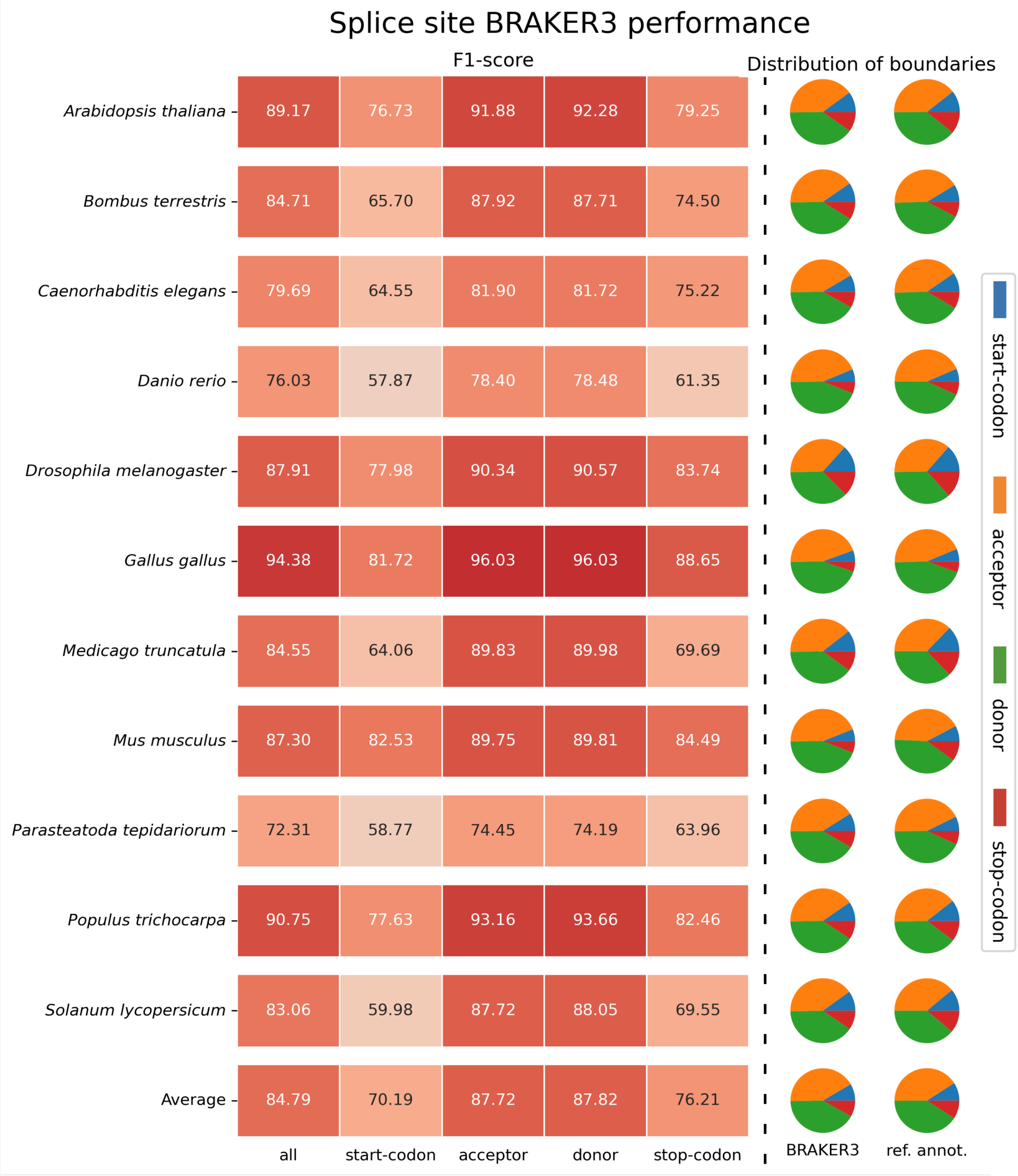


Supplemental Figure S5: F1-score for predicting the signal at the boundaries of coding regions, start- and stop codons and acceptor/donor splice sites, downstream/upstream of an intron, respectively.

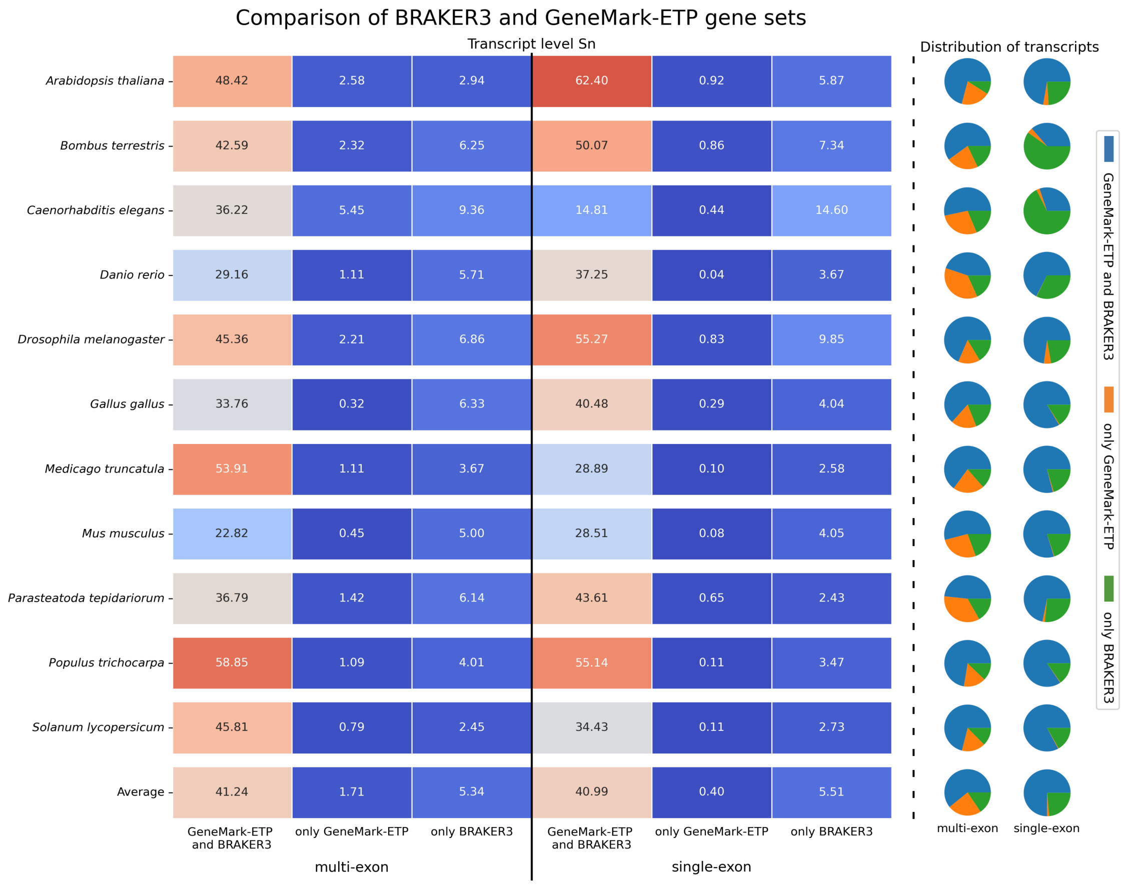


Supplemental Figure S6: The left side of the heat map refers to spliced genes, the right side to unspliced genes. The cells show the percentage of reference transcripts that are correctly predicted by both BRAKER3 and GeneMark-ETP (left columns), by GeneMark-ETP only (middle columns) and by BRAKER3 only (right columns). The pie charts to the right visualize the relative sizes of the predicted transcript sets regardless of whether they are correct.

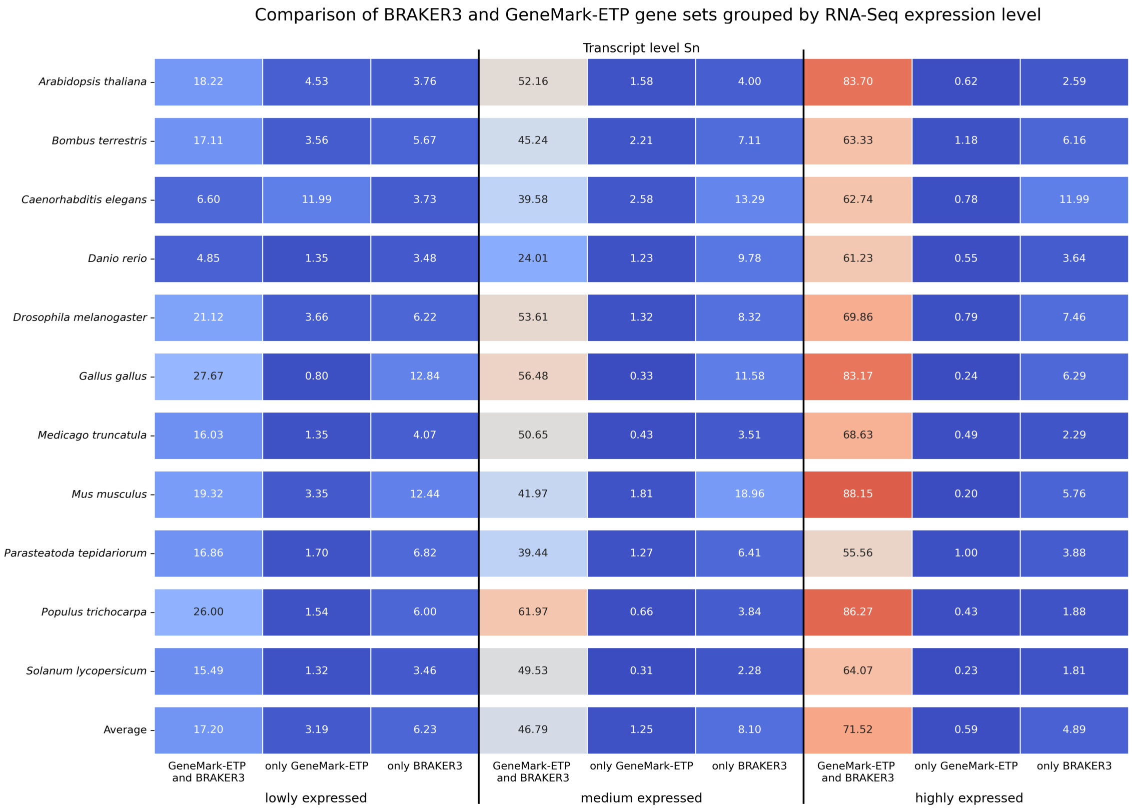


Supplemental Figure S7: The heat map shows the percentage of reference transcripts, in the first, second and third expression tercile, that are correctly identified by BRAKER3 and GeneMark-ETP or by either program only.

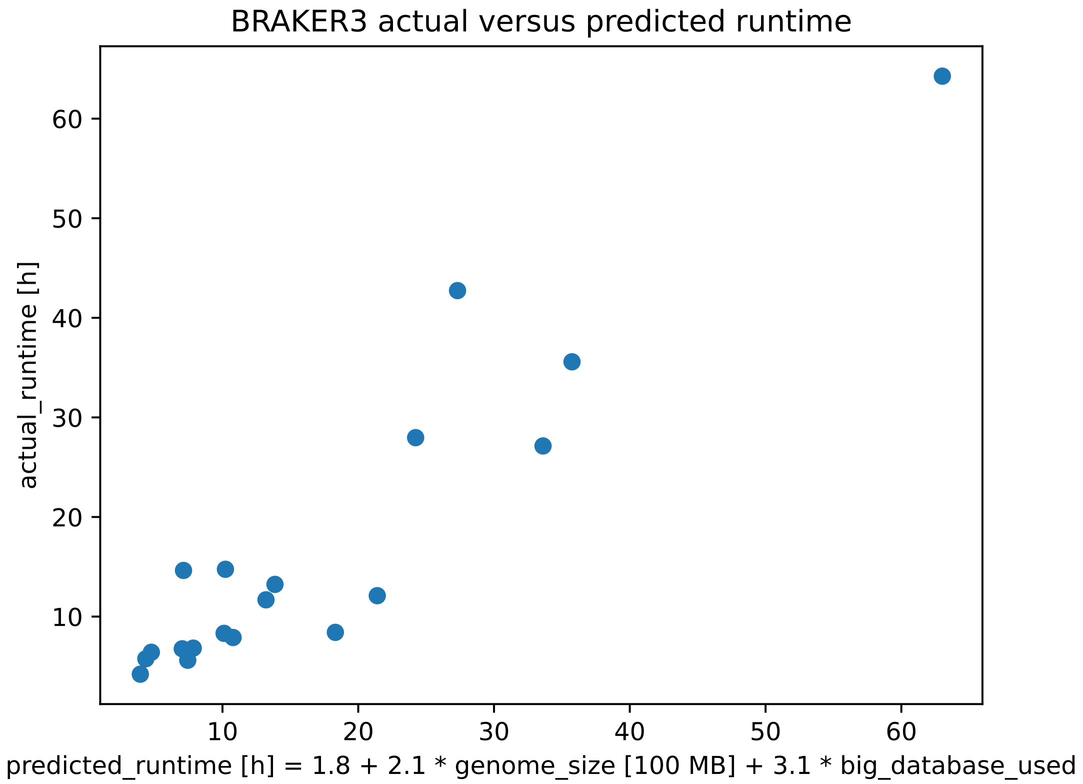


Supplemental Figure S8: Actual runtime versus the runtime predicted for 19 whole genomes. The regression to predict the runtime ($R^2 = 0.87$) considered only the size of the genome and whether an OrthoDB partition was used (`big_database_used`=1) or only the proteomes of a few closely related genomes were used (`big_database_used`=0).

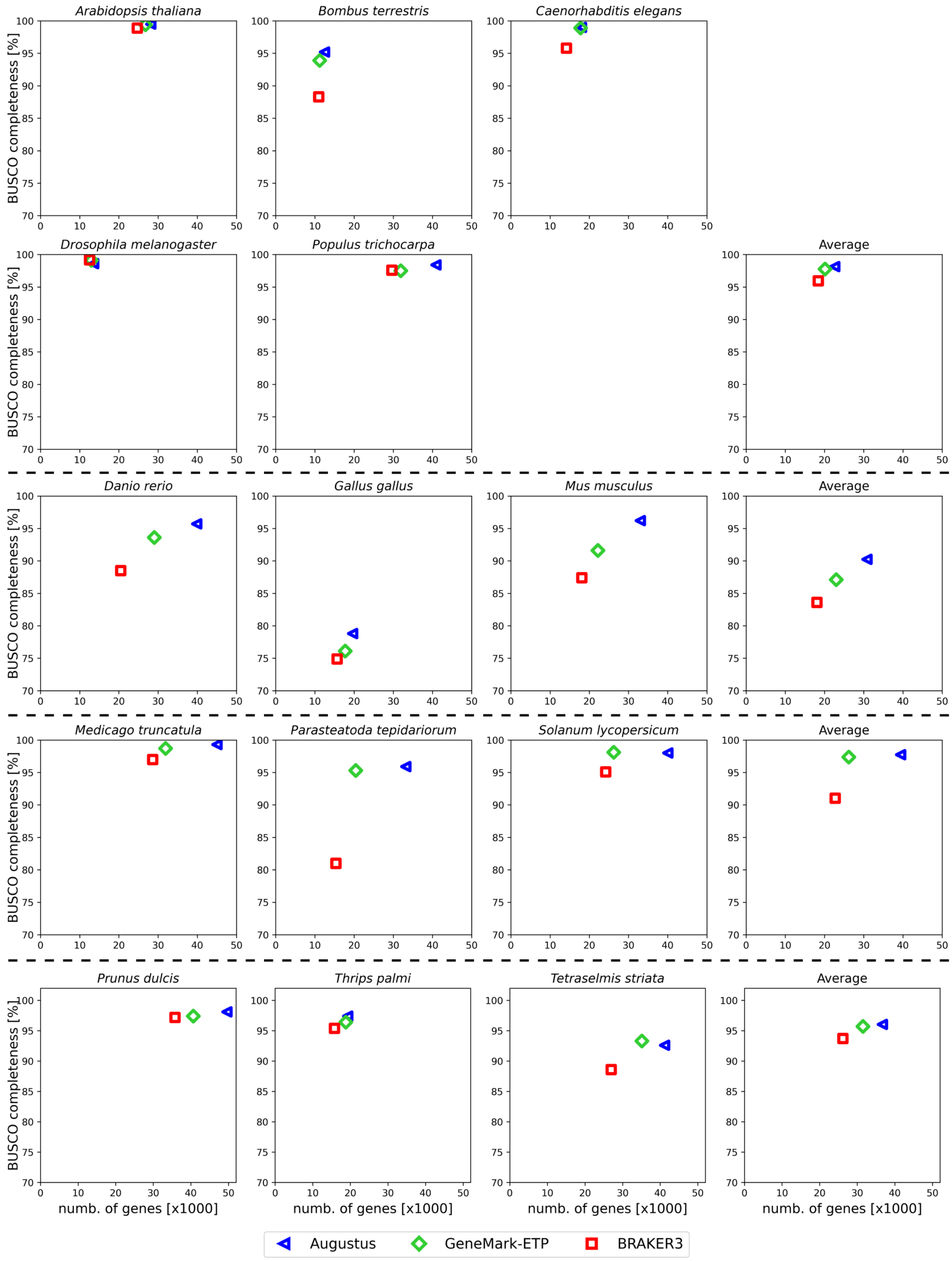


Supplemental Figure S9: Proteome completeness versus the number of genes. The horizontal axis shows the total gene count, after alternative transcripts were grouped into genes. The vertical axis shows the BUSCO completeness percentage (single-copy or duplicated) for the respective gene sets. The AUGUSTUS and GeneMark-ETP gene sets were taken from the output of BRAKER3.

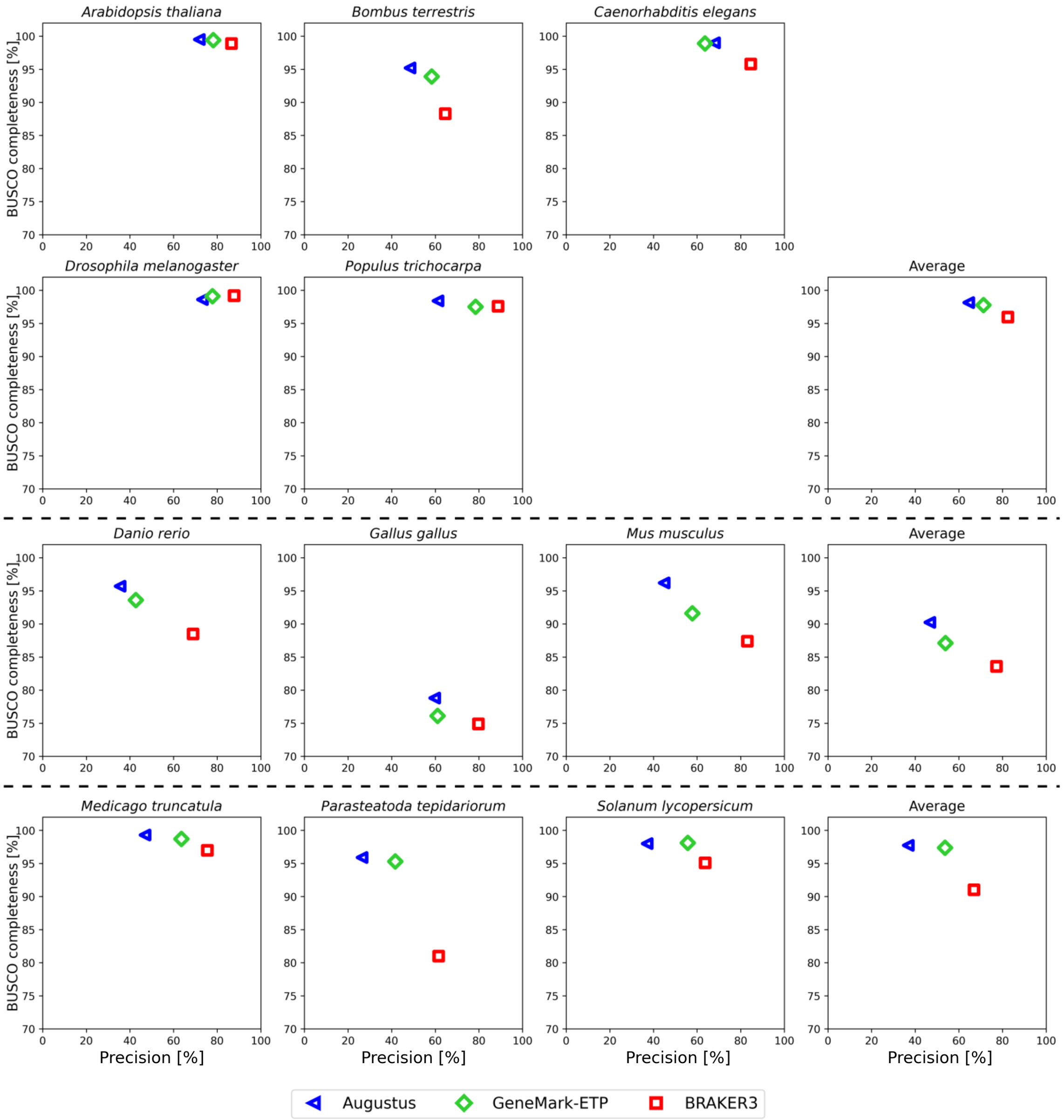


Supplemental Figure S10: Proteome completeness versus gene-level precision. The data is as in Supplemental Figure S9, except that the horizontal axis shows the percentage of predicted genes that identically share a transcript with the reference annotation.

#### 1.2 Supplemental Tables

| Species | Annotation version | Size (Mbp) | #Genes | #Transcripts | #CDS | # Sequences in protein database | | |
|---|---|---|---|---|---|---|---|---|
| | | | | | | *species excluded* | *order excluded* | *close relatives included* |
| *Arabidopsis thaliana* | TAIR Araport 11 (Jun 2016) | 119 | 27,445 | 48,149 | 156,731 | 5,283,100 | 4,825,128 | 306,719 |
| *Bombus terrestris* | NCBI Annotation Release 102 (Apr 2017) | 249 | 10,581 | 22,091 | 78,337 | 4,297,173 | 3,416,393 | 180,811 |
| *Caenorhabditis elegans* | WormBase WS271 (May 2019) | 100 | 20,172 | 33,624 | 130,885 | 15,237,008 | 15,145,380 | 115,553 |
| *Danio rerio* | Ensembl GRCz11.96 (May 2019) | 1,345 | 25,611 | 42,934 | 262,325 | 9,779,764 | 9,468,332 | 760,754 |
| *Drosophila melanogaster* | FlyBase R6.18 (Jun 2019) | 138 | 13,930 | 30,561 | 62,841 | 4,293,925 | 3,029,616 | 116,493 |
| *Gallus gallus* | Ensembl GRCg6a.105 (March 2018) | 1,050 | 17,279 | 38,534 | 192,095 | 9,787,814 | 9,690,812 | 220,641 |
| *Medicago truncatula* | MtrunA17r5.0-ANR-EGN-r1.6 (Feb 2019) | 420 | 44,464 | 44,464 | 174,281 | 5,278,627 | 4,680,430 | 134,774 |
| *Mus musculus* | GENCODE GRCm39 version M28 | 2,723 | 22,405 | 58,318 | 243,366 | 9,782,804 | 9,059,968 | 510,476 |
| *Parasteatoda tepidariorum* | NCBI Annotation Release 101 (May 2017) | 1,445 | 18,602 | 27,516 | 143,792 | 4,287,893 | 4,240,214 | 1,163,197 |
| *Populus trichocarpa* | JGI Ptrichocarpa_533_v4.1 (Nov 2019) | 389 | 34,488 | 52,085 | 187,170 | 5,278,879 | 5,112,927 | 199,662 |
| *Solanum lycopersicum* | Consortium ITAG4.0 (May 2019) | 773 | 33,562 | 33,562 | 152,352 | 5,284,979 | 4,823,899 | 149,444 |

Supplemental Table S1: Summary of genomes, annotations and protein databases used for performance evaluation. Data extracted from Table 4 in Brůna et al. (2023b) and computed from raw data of Brůna et al. (2021, 2023b). Genome versions, repeat masking and annotation processing are documented at `https://github.com/gatech-genemark/EukSpecies-BRAKER2` and at `https://github.com/gatech-genemark/GeneMark-ETP-exp`. The protein databases were generated with orthodb-clades from `https://github.com/tomasbruna/orthodb-clades`.

| **Species** | **Reference Protein File** |
|---|---|
| Arabidopsis thaliana | |
| Arabidopsis lyrata subsp. lyrata | GCF_000004255.2_v.1.0_protein.faa.gz |
| Arabidopsis thaliana x Arabidopsis arenosa | GCA_019202795.1_ASM1920279v1_protein.faa.gz |
| Camelina sativa | GCF_000633955.1_Cs_protein.faa.gz |
| Arabidopsis suecica | GCA_019202805.1_ASM1920280v1_protein.faa.gz |
| Capsella rubella | GCF_000375325.1_Caprub1_0_protein.faa.gz |
| Bombus terrestris | |
| Bombus vancouverensis nearcticus | GCF_011952275.1_Bvanc_JDL1245_protein.faa.gz |
| Bombus huntii | GCF_024542735.1_iyBomHunt1.1_protein.faa.gz |
| Bombus affinis | GCF_024516045.1_iyBomAffi1.2_protein.faa.gz |
| Bombus pyrosoma | GCF_014825855.1_ASM1482585v1_protein.faa.gz |
| Bombus vosnesenskii | GCF_011952255.1_Bvos_JDL3184-5_v1.1_protein.faa.gz |
| Bombus bifarius | GCF_011952205.1_Bbif_JDL3187_protein.faa.gz |
| Bombus impatiens | GCF_000188095.3_BIMP_2.2_protein.faa.gz |
| Caenorhabditis elegans | |
| Caenorhabditis auriculariae | GCA_904845305.1_CAUJ_protein.faa.gz |
| Caenorhabditis bovis | GCA_902829315.1_CBOVIS_v1.1_protein.faa.gz |
| Caenorhabditis brenneri | GCA_000143925.2_C_brenneri-6.0.1b_protein.faa.gz |
| Caenorhabditis briggsae | GCF_000004555.2_CB4_protein.faa.gz |
| Caenorhabditis remanei | GCF_000149515.1_ASM14951v1_protein.faa.gz |
| Danio rerio | |
| Cyprinus carpio | GCF_018340385.1_ASM1834038v1_protein.faa.gz |
| Carassius auratus | GCF_003368295.1_ASM336829v1_protein.faa.gz |
| Puntigrus tetrazona | GCF_018831695.1_ASM1883169v1_protein.faa.gz |
| Sinocyclocheilus rhinocerous | GCF_001515625.1_SAMN03320098_v1.1_protein.faa.gz |
| Sinocyclocheilus anshuiensis | GCF_001515605.1_SAMN03320099.WGS_v1.1_protein.faa.gz |
| Onychostoma macrolepis | GCA_012432095.1_ASM1243209v1_protein.faa.gz |
| Carassius gibelio | GCF_023724105.1_carGib1.2-hapl.c_protein.faa.gz |
| Pimephales promelas | GCF_016745375.1_EPA_FHM_2.0_protein.faa.gz |
| Labeo rohita | GCF_022985175.1_IGBB_LRoh.1.0_protein.faa.gz |
| Megalobrama amblycephala | GCF_018812025.1_ASM1881202v1_protein.faa.gz |
| Sinocyclocheilus grahami | GCF_001515645.1_SAMN03320097.WGS_v1.1_protein.faa.gz |
| Ctenopharyngodon idella | GCF_019924925.1_HZGC01_protein.faa.gz |
| Drosophila melanogaster | |
| Drosophila ananassae | GCF_017639315.1_ASM1763931v2_protein.faa.gz |
| Drosophila grimshawi | GCF_018153295.1_ASM1815329v1_protein.faa.gz |
| Drosophila pseudoobscura | GCF_009870125.1_UCI_Dpse_MV25_protein.faa.gz |
| Drosophila virilis | GCF_003285735.1_DvirRS2_protein.faa.gz |
| Drosophila willistoni | GCF_018902025.1_UCI_dwil_1.1_protein.faa.gz |
| Gallus gallus | |
| Lagopus muta | GCF_023343835.1_bLagMut1_primary_protein.faa.gz |
| Tympanuchus pallidicinctus | GCF_026119805.1_pur_lepc_1.0_protein.faa.gz |
| Lagopus leucura | GCF_019238085.1_USGS_WTPT01_protein.faa.gz |
| Centrocercus urophasianus | GCF_019232065.1_USGS_Curo_1.0_protein.faa.gz |
| Centrocercus urophasianus | GCF_019232065.1_USGS_Curo_1.0_protein.faa.gz |
| Coturnix japonica | GCF_001577835.2_Coturnix_japonica_2.1_protein.faa.gz |

| | |
|---|---|
| Meleagris gallopavo | GCF_000146605.3_Turkey_5.1_protein.faa.gz |
| Medicago truncatula | |
| Trifolium pratense | GCF_020283565.1_ARS_RC_1.1_protein.faa.gz |
| Pisum sativum | GCF_024323335.1_CAAS_Psat_ZW6_1.0_protein.faa.gz |
| Cicer arietinum | GCF_000331145.1_ASM33114v1_protein.faa.gz |
| Mus musculus | |
| Arvicanthis niloticus | GCF_011762505.1_mArvNil1.pat.X_protein.faa.gz |
| Grammomys surdaster | GCF_004785775.1_NIH_TR_1.0_protein.faa.gz |
| Mastomys coucha | GCF_008632895.1_UCSF_Mcou_1_protein.faa.gz |
| Mus pahari | GCF_900095145.1_PAHARI_EIJ_v1.1_protein.faa.gz |
| Apodemus sylvaticus | GCF_947179515.1_mApoSyl1.1_protein.faa.gz |
| Mus caroli | GCF_900094665.1_CAROLI_EIJ_v1.1_protein.faa.gz |
| Rattus rattus | GCF_011064425.1_Rrattus_CSIRO_v1_protein.faa.gz |
| Rattus norvegicus | GCF_015227675.2_mRatBN7.2_protein.faa.gz |
| Homo sapiens | GCF_000001405.40_GRCh38.p14_protein.faa.gz |
| Parasteatoda tepidariorum | |
| Trichonephila inaurata | GCA_019973955.1_Tnin_1.0_protein.faa.gz |
| Caerostris extrusa | GCA_021605095.1_Cext_1.0_protein.faa.gz |
| Caerostris darwini | GCA_021605075.1_Cdar_1.0_protein.faa.gz |
| Oedothorax gibbosus | GCA_019343175.1_Ogib_1.0_protein.faa.gz |
| Trichonephila clavata | GCA_019973975.1_Tnct_1.0_protein.faa.gz |
| Trichonephila clavipes | GCA_019973935.1_Tncv_1.0_protein.faa.gz |
| Araneus ventricosus | GCA_013235015.1_Ave_3.0_protein.faa.gz |
| Nephila pilipes | GCA_019974015.1_Npil_1.0_protein.faa.gz |
| Populus trichocarpa | |
| Populus tomentosa | GCA_018804465.1_PTv2_protein.faa.gz |
| Populus euphratica | GCF_000495115.1_PopEup_1.0_protein.faa.gz |
| Populus alba | GCF_005239225.1_ASM523922v1_protein.faa.gz |
| Populus deltoides | GCA_015852605.2_ASM1585260v2_protein.faa.gz |
| Solanum lycopersicum | |
| Solanum stenotomum | GCF_019186545.1_ASM1918654v1_protein.faa.gz |
| Solanum tuberosum | GCF_000226075.1_SolTub_3.0_protein.faa.gz |
| Solanum verrucosum | GCF_900185275.1_falcon-dt-bn_protein.faa.gz |
| Solanum pennellii | GCF_001406875.1_SPENNV200_protein.faa.gz |

Supplemental Table S2: Donor proteins used for each species for the close relative included protein set.

| Species | OrthoDB database | # BUSCO |
|---|---|---:|
| *Arabidopsis thaliana* | eudicots_odb10 | 2,326 |
| *Bombus terrestris* | hymenoptera_odb10 | 5,991 |
| *Caenorhabditis elegans* | nematoda_odb10 | 3,131 |
| *Danio rerio* | actinopterygii_odb10 | 3,640 |
| *Drosophila melanogaster* | arthropoda_odb10 | 1,013 |
| *Gallus gallus* | mammalia_odb10 | 9,226 |
| *Medicago truncatula* | fabales_odb10 | 5,366 |
| *Mus musculus* | mammalia_odb10 | 9,226 |
| *Parasteatoda tepidariorum* | arachnida_odb10 | 2,934 |
| *Populus trichocarpa* | eudicots_odb10 | 2,326 |
| *Prunus dulcis* | viridiplantae_odb10 | 425 |
| *Solanum lycopersicum* | solanales_odb10 | 5,950 |
| *Thrips palmi* | insecta_odb10 | 1,367 |
| *Tetraselmis striata* | chlorophyta_odb10 | 1,519 |

Supplemental Table S3: Summary of OrthoDB databases and number of BUSCOs used for BUSCO evaluation on each test species.

| | Exon | | Gene | | Transcript | | Exon | | Gene | | Transcript | |
|---|---|---|---|---|---|---|---|---|---|---|---|---|
| | Sn | Prec | Sn | Prec | Sn | Prec | Sn | Prec | Sn | Prec | Sn | Prec |
| | *Arabidopsis thaliana* | | | | | | *Bombus terrestris* | | | | | |
| BRAKER1 | 78.20 | 80.83 | 58.37 | 58.73 | 39.97 | 56.75 | 78.86 | 69.77 | 42.02 | 27.45 | 28.72 | 25.67 |
| BRAKER2 | 80.91 | 87.22 | 71.96 | 68.58 | 48.48 | 66.33 | 74.06 | 77.04 | 45.38 | 30.81 | 28.47 | 29.85 |
| TSEBRA | 71.02 | 93.68 | 70.92 | 84.38 | 49.26 | 75.30 | 65.61 | 79.95 | 56.87 | 44.26 | 39.24 | 38.17 |
| GeneMark-ETP | **82.71** | 89.67 | 76.59 | 78.22 | 53.11 | 76.76 | **81.10** | 85.64 | 62.26 | 58.36 | 45.16 | 54.20 |
| BRAKER3 | 78.64 | **95.81** | **77.90** | **86.53** | **54.25** | **82.30** | 74.58 | **89.64** | **66.82** | **64.51** | **49.19** | **56.66** |
| | *Caenorhabditis elegans* | | | | | | *Danio rerio* | | | | | |
| BRAKER1 | **82.00** | 87.55 | 57.58 | 62.28 | 42.87 | 59.15 | 78.00 | 69.80 | 40.35 | 23.48 | 25.14 | 22.36 |
| BRAKER2 | 74.13 | 87.97 | 47.76 | 54.30 | 33.87 | 53.11 | 76.14 | 68.71 | 41.30 | 20.12 | 25.11 | 19.37 |
| TSEBRA | 61.53 | 93.80 | 56.09 | 81.63 | 42.51 | 72.08 | 47.19 | 83.78 | 43.84 | 39.39 | 27.53 | 34.86 |
| GeneMark-ETP | 81.09 | 88.75 | 56.13 | 63.64 | 40.82 | 62.78 | **78.12** | 77.28 | 49.20 | 42.72 | 30.68 | 41.84 |
| BRAKER3 | 65.89 | **95.07** | **59.36** | **84.54** | **45.06** | **76.31** | 61.89 | **91.95** | **54.58** | **68.95** | **34.43** | **60.37** |
| | *Drosophila melanogaster* | | | | | | *Gallus gallus* | | | | | |
| BRAKER1 | 76.81 | 76.77 | 59.58 | 58.69 | 39.40 | 55.26 | 63.28 | 46.92 | 5.96 | 2.97 | 5.09 | 2.88 |
| BRAKER2 | 71.03 | 83.23 | 60.18 | 59.77 | 37.80 | 58.16 | 33.49 | 59.35 | 25.28 | 16.46 | 21.38 | 16.22 |
| TSEBRA | 65.72 | 89.99 | 69.18 | 79.69 | 46.26 | 70.45 | 26.85 | 77.34 | 25.61 | 27.11 | 21.77 | 26.03 |
| GeneMark-ETP | **78.11** | 89.78 | 72.68 | 77.85 | 48.77 | 75.61 | **94.20** | 87.49 | 76.41 | 61.06 | 68.87 | 58.24 |
| BRAKER3 | 77.13 | **94.46** | **78.71** | **87.73** | **54.03** | **79.96** | 93.57 | **94.79** | **84.11** | **79.73** | **75.25** | **70.05** |
| | *Medicago truncatula* | | | | | | *Mus musculus* | | | | | |
| BRAKER1 | 76.77 | 65.93 | 36.95 | 37.59 | 36.95 | 35.78 | 82.22 | 66.24 | 28.01 | 13.09 | 28.00 | 12.70 |
| BRAKER2 | 78.71 | 71.11 | 44.91 | 43.77 | 44.91 | 42.16 | 61.45 | 71.56 | 37.47 | 20.84 | 37.46 | 20.44 |
| TSEBRA | 66.50 | 85.33 | 46.01 | 62.73 | 46.01 | 53.97 | 51.58 | 79.74 | 42.11 | 30.90 | 42.10 | 27.99 |
| GeneMark-ETP | **79.32** | 79.62 | 45.86 | 63.62 | 45.86 | 59.80 | **89.50** | 88.14 | 67.47 | 57.85 | 67.47 | 56.05 |
| BRAKER3 | 75.07 | **90.31** | **46.96** | **75.47** | **46.96** | **67.94** | 81.81 | **96.73** | **75.84** | **83.05** | **75.83** | **73.11** |
| | *Parasteatoda tepidariorum* | | | | | | *Populus trichocarpa* | | | | | |
| BRAKER1 | **77.49** | 63.48 | 41.65 | 22.80 | 34.25 | 21.45 | 81.13 | 72.11 | 50.56 | 44.31 | 40.52 | 42.29 |
| BRAKER2 | 67.96 | 58.10 | 29.75 | 14.17 | 23.37 | 13.74 | 84.99 | 80.47 | 68.69 | 58.03 | 53.84 | 56.19 |
| TSEBRA | 48.34 | 73.59 | 44.65 | 31.98 | 37.08 | 29.11 | 74.72 | 90.31 | 68.66 | 76.85 | 55.30 | 66.53 |
| GeneMark-ETP | 76.52 | 72.39 | 45.56 | 41.66 | 38.99 | 40.97 | **86.13** | 88.69 | 72.88 | 78.47 | 58.95 | 74.49 |
| BRAKER3 | 57.81 | **87.61** | **47.16** | **61.51** | **40.69** | **55.47** | 83.32 | **95.10** | **74.90** | **88.76** | **60.68** | **81.63** |
| | *Solanum lycopersicum* | | | | | | **Average** | | | | | |
| BRAKER1 | 75.24 | 62.44 | 33.09 | 29.39 | 33.09 | 27.62 | 77.27 | 69.26 | 41.28 | 34.62 | 32.18 | 32.90 |
| BRAKER2 | 76.90 | 66.89 | 41.40 | 33.70 | 41.40 | 32.46 | 70.89 | 73.79 | 46.73 | 38.23 | 36.01 | 37.09 |
| TSEBRA | 68.74 | 79.93 | 43.94 | 50.13 | 43.94 | 42.17 | 58.89 | 84.31 | 51.63 | 55.37 | 41.00 | 48.79 |
| GeneMark-ETP | **77.68** | 80.15 | 43.65 | 55.75 | 43.65 | 51.25 | **82.23** | 84.33 | 60.79 | 61.75 | 49.30 | 59.27 |
| BRAKER3 | 74.05 | **86.85** | **44.46** | **63.66** | **44.46** | **56.51** | 74.89 | **92.57** | **64.62** | **76.77** | **52.80** | **69.12** |

Supplemental Table S4: Sensitivity (Sn) and precision (Prec) for a protein database in which proteins of the same *order* as the target species were *excluded*. The last subtable shows the respective averages for the 11 different species. The highest number in each column is indicated in bold text. Inputs were for each species a genome assembly, short-read RNA-seq libraries, and a protein database (respective **order excluded**).

| | Exon | Gene | Transcript | Exon | Gene | Transcript | Exon | Gene | Transcript | Exon | Gene | Transcript |
|---|---|---|---|---|---|---|---|---|---|---|---|---|
| | *Arabidopsis thaliana* | | | *Bombus terrestris* | | | *Caenorhabditis elegans* | | | *Danio rerio* | | |
| BRAKER1 | 79.49 | 58.55 | 46.90 | 74.04 | 33.21 | 27.11 | 84.68 | 59.84 | 49.71 | 73.67 | 29.69 | 23.67 |
| BRAKER2 | 83.95 | 70.23 | 56.02 | 75.52 | 36.70 | 29.14 | 80.46 | 50.82 | 41.36 | 72.23 | 27.06 | 21.87 |
| TSEBRA | 80.79 | 77.07 | 59.56 | 72.07 | 49.78 | 38.70 | 74.31 | 66.49 | 53.48 | 60.37 | 41.50 | 30.76 |
| GeneMark-ETP | 86.05 | 77.40 | 62.78 | **83.31** | 60.25 | 49.27 | **84.75** | 59.65 | 49.47 | **77.70** | 45.73 | 35.40 |
| BRAKER3 | **86.38** | **81.99** | **65.39** | 81.42 | **65.64** | **52.66** | 77.84 | **69.75** | **56.66** | 73.98 | **60.93** | **43.85** |
| | *Drosophila melanogaster* | | | *Gallus gallus* | | | *Medicago truncatula* | | | *Mus musculus* | | |
| BRAKER1 | 76.79 | 59.13 | 46.00 | 53.89 | 3.96 | 3.68 | 70.94 | 37.27 | 36.36 | 73.37 | 17.84 | 17.47 |
| BRAKER2 | 76.65 | 59.97 | 45.82 | 42.82 | 19.94 | 18.45 | 74.72 | 44.33 | 43.49 | 66.12 | 26.78 | 26.45 |
| TSEBRA | 75.96 | 74.06 | 55.85 | 39.86 | 26.34 | 23.71 | 74.75 | 53.08 | 49.67 | 62.64 | 35.64 | 33.62 |
| GeneMark-ETP | 83.54 | 75.18 | 59.29 | 90.72 | 67.88 | 63.11 | 79.47 | 53.30 | 51.91 | **88.81** | 62.29 | 61.23 |
| BRAKER3 | **84.92** | **82.98** | **64.49** | **94.18** | **81.86** | **72.56** | **81.99** | **57.90** | **55.53** | 88.65 | **79.28** | **74.45** |
| | *Parasteatoda tepidariorum* | | | *Populus trichocarpa* | | | *Solanum lycopersicum* | | | **Average** | | |
| BRAKER1 | 69.79 | 29.47 | 26.38 | 76.35 | 47.23 | 41.39 | 68.24 | 31.13 | 30.11 | 73.05 | 37.66 | 32.54 |
| BRAKER2 | 62.64 | 19.20 | 17.31 | 82.67 | 62.91 | 54.99 | 71.55 | 37.16 | 36.39 | 72.31 | 42.06 | 36.54 |
| TSEBRA | 58.35 | 37.27 | 32.62 | 81.78 | 72.52 | 60.40 | 73.91 | 46.83 | 43.04 | 69.35 | 53.43 | 44.56 |
| GeneMark-ETP | **74.40** | 43.52 | 39.96 | 87.39 | 75.57 | 65.82 | 78.90 | 48.96 | 47.15 | **83.26** | 61.26 | 53.83 |
| BRAKER3 | 69.66 | **53.39** | **46.94** | **88.82** | **81.24** | **69.61** | **79.94** | **52.36** | **49.77** | 82.80 | **70.17** | **59.87** |

Supplemental Table S5: F1-scores of pipelines obtaining short-read RNA-seq libraries, and a protein database (respective **order excluded**) as input. The subtable on the bottom right shows the averages for the 11 different species. The highest number in each column is indicated in bold text.

| | Exon | | Gene | | Transcript | | Exon | | Gene | | Transcript | |
|---|---|---|---|---|---|---|---|---|---|---|---|---|
| | Sn | Prec | Sn | Prec | Sn | Prec | Sn | Prec | Sn | Prec | Sn | Prec |
| | | | *Arabidopsis thaliana* | | | | | | *Bombus terrestris* | | | |
| BRAKER1 | 78.20 | 80.83 | 58.37 | 58.73 | 39.97 | 56.75 | 78.86 | 69.77 | 42.02 | 27.45 | 28.72 | 25.67 |
| BRAKER2 | 83.20 | 86.83 | 79.18 | 72.88 | 53.81 | 69.00 | 81.29 | 75.63 | 61.9 | 35.57 | 39.90 | 33.96 |
| TSEBRA | 81.58 | 92.89 | 81.69 | 83.74 | 56.83 | 74.89 | 72.38 | 82.45 | 64.17 | 43.21 | 42.36 | 41.68 |
| GeneMark-ETP | **83.45** | 89.71 | 78.83 | 79.25 | 54.89 | 77.65 | **83.37** | 86.25 | 66.87 | 61.38 | 49.38 | 57.21 |
| BRAKER3 | 81.72 | **95.80** | **81.8** | **87.23** | **57.01** | **83.85** | 78.56 | **90.10** | **72.51** | **66.34** | **52.53** | **60.08** |
| | | | *Caenorhabditis elegans* | | | | | | *Danio rerio* | | | |
| BRAKER1 | 82.00 | 87.55 | 57.58 | 62.28 | 42.87 | 59.15 | 78.00 | 69.80 | 40.35 | 23.48 | 25.14 | 22.36 |
| BRAKER2 | 83.41 | 90.53 | 66.22 | 67.44 | 47.34 | 63.67 | 77.46 | 66.77 | 44.71 | 20.21 | 27.21 | 19.25 |
| TSEBRA | 76.22 | 93.85 | **69.04** | 82.25 | **51.92** | 71.96 | 60.70 | 82.37 | 53.19 | 39.6 | 33.34 | 34.8 |
| GeneMark-ETP | **83.84** | 90.07 | 63.98 | 69.45 | 47.73 | 67.38 | **78.71** | 76.58 | 50.27 | 41.58 | 31.34 | 40.77 |
| BRAKER3 | 75.25 | **95.45** | 68.22 | **85.54** | 51.86 | **77.64** | 65.15 | **90.79** | **56.91** | **66.60** | **35.83** | **58.23** |
| | | | *Drosophila melanogaster* | | | | | | *Gallus gallus* | | | |
| BRAKER1 | 76.81 | 76.77 | 59.58 | 58.69 | 39.40 | 55.26 | 63.28 | 46.92 | 5.96 | 2.97 | 5.09 | 2.88 |
| BRAKER2 | 79.64 | 88.45 | 77.78 | 73.76 | 49.63 | 70.36 | 36.16 | 61.33 | 28.56 | 19.31 | 24.25 | 18.93 |
| TSEBRA | 78.14 | 91.68 | 82.20 | 84.29 | 55.09 | 74.08 | 32.87 | 80.64 | 31.22 | 32.28 | 26.61 | 30.99 |
| GeneMark-ETP | 82.49 | 91.71 | 80.24 | 83.59 | 55.84 | 79.91 | **94.14** | 86.96 | 75.66 | 59.50 | 68.20 | 57.26 |
| BRAKER3 | **82.88** | **95.75** | **85.67** | **89.89** | **58.66** | **83.69** | 93.92 | **94.52** | **84.87** | **79.57** | **75.95** | **70.07** |
| | | | *Medicago truncatula* | | | | | | *Mus musculus* | | | |
| BRAKER1 | 76.77 | 65.93 | 36.95 | 37.59 | 36.95 | 35.78 | 82.22 | 66.24 | 28.01 | 13.09 | 28.00 | 12.70 |
| BRAKER2 | 79.84 | 71.47 | 47.09 | 46.56 | 47.09 | 43.98 | 61.70 | 71.34 | 40.04 | 22.10 | 40.03 | 21.54 |
| TSEBRA | 75.63 | 84.84 | 48.63 | 57.96 | 48.63 | 55.16 | 61.87 | 81.82 | 51.82 | 34.27 | 51.81 | 31.10 |
| GeneMark-ETP | **80.16** | 77.53 | 47.85 | 61.38 | 47.85 | 57.67 | **90.15** | 87.52 | 68.34 | 55.90 | 68.34 | 54.38 |
| BRAKER3 | 76.89 | **88.98** | **49.62** | **73.14** | **49.62** | **66.57** | 85.37 | **96.40** | **79.62** | **81.50** | **79.61** | **71.91** |
| | | | *Parasteatoda tepidariorum* | | | | | | *Populus trichocarpa* | | | |
| BRAKER1 | **77.49** | 63.48 | 41.65 | 22.80 | 34.25 | 21.45 | 81.13 | 72.11 | 50.56 | 44.31 | 40.52 | 42.29 |
| BRAKER2 | 69.11 | 58.97 | 32.24 | 15.06 | 25.33 | 14.46 | 86.45 | 80.58 | 73.36 | 61.11 | 57.90 | 58.52 |
| TSEBRA | 50.45 | 72.84 | 45.54 | 30.48 | 37.60 | 28.14 | 84.72 | 90.13 | **77.75** | 77.36 | 62.71 | 67.22 |
| GeneMark-ETP | 77.14 | 71.99 | 47.12 | 41.42 | 40.78 | 40.74 | **86.65** | 88.07 | 74.30 | 77.65 | 60.17 | 73.73 |
| BRAKER3 | 58.53 | **86.70** | **48.64** | **59.63** | **42.14** | **54.05** | 85.08 | **94.99** | 77.75 | **88.31** | **63.08** | **82.00** |
| | | | *Solanum lycopersicum* | | | | | | **Average** | | | |
| BRAKER1 | 75.24 | 62.44 | 33.09 | 29.39 | 33.09 | 27.62 | 77.27 | 69.26 | 41.28 | 34.62 | 32.18 | 32.90 |
| BRAKER2 | 78.43 | 60.74 | 44.59 | 31.55 | 44.59 | 29.25 | 74.24 | 73.88 | 54.15 | 42.32 | 41.55 | 40.27 |
| TSEBRA | 75.69 | 77.47 | **48.46** | 47.21 | **48.46** | 40.09 | 68.20 | 84.63 | 59.43 | 55.70 | 46.85 | 50.01 |
| GeneMark-ETP | **78.59** | 76.20 | 45.37 | 51.99 | 45.37 | 47.82 | **83.52** | 83.87 | 63.53 | 62.1 | 51.81 | 59.50 |
| BRAKER3 | 75.87 | **85.37** | 46.90 | **60.37** | 46.90 | **54.72** | 78.11 | **92.26** | **68.41** | **76.19** | **55.74** | **69.35** |

Supplemental Table S6: Sensitivity (Sn) and precision (Prec) for a protein database in which only proteins from the target species were *excluded*. The last subtable shows the respective averages for the 11 different species. The highest number in each column is indicated in bold text. Inputs were for each species a genome assembly, short-read RNA-seq libraries, and a protein database (respective **species excluded**).

| | Exon | Gene | Transcript | Exon | Gene | Transcript | Exon | Gene | Transcript | Exon | Gene | Transcript |
|---|---|---|---|---|---|---|---|---|---|---|---|---|
| | *Arabidopsis thaliana* | | | *Bombus terrestris* | | | *Caenorhabditis elegans* | | | *Danio rerio* | | |
| BRAKER1 | 79.49 | 58.55 | 46.90 | 74.04 | 33.21 | 27.11 | 84.68 | 59.84 | 49.71 | 73.67 | 29.69 | 23.67 |
| BRAKER2 | 84.98 | 75.90 | 60.47 | 78.36 | 45.18 | 36.69 | 86.82 | 66.82 | 54.30 | 71.72 | 27.84 | 22.55 |
| TSEBRA | 86.87 | 82.70 | 64.62 | 77.09 | 51.64 | 42.02 | 84.12 | 75.07 | 60.32 | 69.89 | 45.40 | 34.05 |
| GeneMark-ETP | 86.47 | 79.04 | 64.32 | **84.79** | 64.01 | 53.01 | **86.84** | 66.60 | 55.88 | **77.63** | 45.51 | 35.44 |
| BRAKER3 | **88.20** | **84.43** | **67.87** | 83.94 | **69.29** | **56.05** | 84.15 | **75.90** | **62.18** | 75.86 | **61.37** | **44.36** |
| | *Drosophila melanogaster* | | | *Gallus gallus* | | | *Medicago truncatula* | | | *Mus musculus* | | |
| BRAKER1 | 76.79 | 59.13 | 46.00 | 53.89 | 3.96 | 3.68 | 70.94 | 37.27 | 36.36 | 73.37 | 17.84 | 17.47 |
| BRAKER2 | 83.81 | 75.72 | 58.20 | 45.50 | 23.04 | 21.26 | 75.42 | 46.82 | 45.48 | 66.17 | 28.48 | 28.01 |
| TSEBRA | 84.37 | 83.23 | 63.19 | 46.70 | 31.74 | 28.63 | 79.97 | 52.89 | 51.69 | 70.46 | 41.26 | 38.87 |
| GeneMark-ETP | 86.86 | 81.88 | 65.74 | 90.41 | 66.61 | 62.25 | 78.82 | 53.78 | 52.30 | 88.82 | 61.50 | 60.57 |
| BRAKER3 | **88.85** | **87.73** | **68.97** | **94.22** | **82.13** | **72.89** | **82.49** | **59.13** | **56.86** | **90.55** | **80.55** | **75.56** |
| | *Parasteatoda tepidariorum* | | | *Populus trichocarpa* | | | *Solanum lycopersicum* | | | **Average** | | |
| BRAKER1 | 69.79 | 29.47 | 26.38 | 76.35 | 47.23 | 41.39 | 68.24 | 31.13 | 30.11 | 73.05 | 37.66 | 32.54 |
| BRAKER2 | 63.64 | 20.53 | 18.41 | 83.41 | 66.68 | 58.21 | 68.46 | 36.95 | 35.33 | 74.06 | 47.51 | 40.90 |
| TSEBRA | 59.61 | 36.52 | 32.19 | 87.34 | 77.55 | 64.89 | 76.57 | 47.83 | 43.88 | 75.54 | 57.50 | 48.38 |
| GeneMark-ETP | **74.48** | 44.09 | 40.76 | 87.35 | 75.94 | 66.26 | 77.38 | 48.45 | 46.56 | 83.69 | 62.81 | 55.39 |
| BRAKER3 | 69.88 | **53.58** | **47.36** | **89.76** | **82.69** | **71.31** | **80.34** | **52.79** | **50.51** | **84.60** | **72.09** | **61.81** |

Supplemental Table S7: F1-scores of pipelines obtaining short-read RNA-seq libraries, and a protein database (respective **species excluded**) as input. The subtable on the bottom right shows the averages for the 11 different species. The highest number in each column is indicated in bold text.

| | Exon | | | Gene | | | Transcript | | |
|---|---|---|---|---|---|---|---|---|---|
| | Sn | Prec | F1 | Sn | Prec | F1 | Sn | Prec | F1 |
| | | | | *Arabidopsis thaliana* | | | | | |
| MAKER2* | 78.05 | 82.02 | 79.99 | 60.57 | 57.69 | 59.09 | 40.72 | 57.69 | 47.74 |
| Funannotate | 82.07 | 93.21 | 87.29 | 75.26 | 79.58 | 77.36 | 50.59 | 79.58 | 61.86 |
| BRAKER3 | **83.03** | **94.38** | **88.34** | **82.93** | **86.27** | **84.57** | **58.70** | **80.78** | **67.99** |
| | | | | *Bombus terrestris* | | | | | |
| MAKER2* | 74.12 | 75.22 | 74.67 | 50.24 | 45.57 | 47.79 | 31.55 | 36.75 | 33.95 |
| Funannotate | 76.23 | 72.72 | 74.43 | 51.73 | 30.80 | 38.61 | 32.11 | 30.85 | 31.47 |
| BRAKER3 | **79.39** | **88.71** | **83.79** | **73.07** | **67.32** | **70.08** | **57.86** | **57.31** | **57.58** |
| | | | | *Caenorhabditis elegans* | | | | | |
| MAKER2* | 73.13 | 76.63 | 74.84 | 41.31 | 41.10 | 41.20 | 29.34 | 38.56 | 33.32 |
| Funannotate | **79.71** | 84.19 | 81.89 | 49.44 | 52.16 | 50.76 | 35.01 | 52.18 | 41.90 |
| BRAKER3 | 76.39 | **95.35** | **84.82** | **68.97** | **85.61** | **76.39** | **52.52** | **77.33** | **62.55** |
| | | | | *Drosophila melanogaster* | | | | | |
| MAKER2* | 75.28 | 73.11 | 74.18 | 61.10 | 52.77 | 56.63 | 38.26 | 52.77 | 44.36 |
| Funannotate | 73.87 | 82.66 | 78.02 | 62.87 | 62.98 | 62.92 | 39.38 | 62.99 | 48.46 |
| BRAKER3 | **81.64** | **95.18** | **87.89** | **83.36** | **90.55** | **86.81** | **59.71** | **81.58** | **68.95** |
| | | | | *Gallus gallus* | | | | | |
| MAKER2* | 85.98 | 78.83 | 82.25 | 49.42 | 38.65 | 43.38 | 41.99 | 32.64 | 36.73 |
| Funannotate | 58.75 | 71.22 | 64.39 | 32.23 | 23.20 | 26.98 | 27.19 | 23.23 | 25.05 |
| BRAKER3 | **93.82** | **93.99** | **93.90** | **85.80** | **78.88** | **82.19** | **78.39** | **66.65** | **72.04** |
| | | | | *Medicago truncatula* | | | | | |
| MAKER2* | 69.63 | 73.87 | 71.69 | 33.48 | 49.12 | 39.82 | 33.48 | 45.67 | 38.64 |
| Funannotate | 74.71 | 63.54 | 68.67 | 35.26 | 37.40 | 36.30 | 35.26 | 37.40 | 36.30 |
| BRAKER3 | **74.76** | **90.14** | **81.73** | **47.86** | **75.25** | **58.51** | **47.86** | **68.22** | **56.25** |
| | | | | *Populus trichocarpa* | | | | | |
| MAKER2* | 74.04 | 71.96 | 72.99 | 41.76 | 42.67 | 42.21 | 32.64 | 38.27 | 35.23 |
| Funannotate | 82.44 | 79.30 | 80.84 | 60.83 | 54.90 | 57.71 | 47.38 | 54.90 | 50.86 |
| BRAKER3 | **84.56** | **92.94** | **88.55** | **76.99** | **85.60** | **81.07** | **63.07** | **76.49** | **69.13** |
| | | | | *Solanum lycopersicum* | | | | | |
| MAKER2* | 74.16 | 45.21 | 56.17 | 33.88 | 18.85 | 24.22 | 33.88 | 18.85 | 24.22 |
| Funannotate | 74.62 | 65.91 | 70.00 | 36.12 | 34.10 | 35.08 | 36.12 | 34.10 | 35.08 |
| BRAKER3 | **74.87** | **85.25** | **79.72** | **45.90** | **61.67** | **52.63** | **45.90** | **53.49** | **49.41** |

Supplemental Table S8: Sensitivity, precision, and F1-score for MAKER2, Funannotate, and BRAKER3 using the **close relatives included** protein databases. *The values for MAKER2 may overestimate the realistic performance on new genomes (see main text).

| | Exon | | | Gene | | | Transcript | | |
|---|---|---|---|---|---|---|---|---|---|
| | Sn | Prec | F1 | Sn | Prec | F1 | Sn | Prec | F1 |
| *Arabidopsis thaliana* | 18.0 | 50.42 | 26.53 | 6.03 | 18.42 | 9.09 | 4.16 | 16.68 | 6.66 |
| *Caenorhabditis elegans* | 28.62 | 50.67 | 36.58 | 8.10 | 12.4 | 9.80 | 5.89 | 11.54 | 7.80 |
| *Danio rerio* | 15.16 | 44.12 | 22.57 | 3.26 | 7.17 | 4.48 | 1.99 | 7.06 | 3.10 |
| *Drosophila melanogaster* | 24.28 | 53.95 | 33.49 | 11.32 | 23.01 | 15.17 | 7.48 | 21.18 | 11.06 |
| *Medicago truncatula* | 17.79 | 48.88 | 26.09 | 3.92 | 16.87 | 6.36 | 3.92 | 15.54 | 6.26 |
| *Populus trichocarpa* | 26.99 | 50.07 | 35.07 | 7.37 | 15.34 | 9.96 | 5.8 | 14.75 | 8.33 |
| *Solanum lycopersicum* | 31.21 | 43.98 | 36.51 | 6.41 | 12.68 | 8.52 | 6.41 | 10.12 | 7.85 |
| **Average** | 23.15 | 48.87 | 30.98 | 6.63 | 15.13 | 9.05 | 5.09 | 13.84 | 7.29 |

Supplemental Table S9: Sensitivity, precision, and F1-score of FINDER for runs using the same input data (genomic sequence + proteins + RNA-seq) as in the experiments from Supplemental Table S4. However, FINDER exited with an error for 4 out of the 11 species tested, and we therefore do not report its performance for those species.

| | Exon | | | Gene | | | Transcript | | |
|---|---|---|---|---|---|---|---|---|---|
| | Sn | Prec | F1 | Sn | Prec | F1 | Sn | Prec | F1 |
| *Arabidopsis thaliana* | 83.4 | 87.70 | 85.50 | 78.78 | 75.18 | 76.94 | 54.18 | 71.77 | 61.75 |
| *Bombus terrestris* | 84.49 | 84.17 | 84.33 | 67.38 | 55.01 | 60.57 | 46.56 | 50.28 | 48.35 |
| *Caenorhabditis elegans* | 84.97 | 91.99 | 88.34 | 70.36 | 74.46 | 72.35 | 51.99 | 70.34 | 59.79 |
| *Danio rerio* | 81.74 | 75.80 | 78.66 | 58.06 | 36.21 | 44.6 | 36.29 | 33.04 | 34.59 |
| *Drosophila melanogaster* | 83.49 | 89.93 | 86.59 | 80.87 | 80.71 | 80.79 | 54.09 | 75.66 | 63.08 |
| *Gallus gallus* | 96.17 | 89.11 | 92.51 | 81.70 | 61.03 | 69.87 | 71.73 | 57.34 | 63.73 |
| *Medicago truncatula* | 81.16 | 69.84 | 75.08 | 48.93 | 46.70 | 47.79 | 48.93 | 43.68 | 46.16 |
| *Mus musculus* | 94.51 | 85.48 | 89.77 | 79.46 | 46.23 | 58.45 | 79.46 | 43.42 | 56.15 |
| *Parasteatoda tepidariorum* | 78.67 | 65.55 | 71.51 | 48.13 | 26.32 | 34.03 | 39.77 | 24.53 | 30.34 |
| *Populus trichocarpa* | 87.70 | 81.99 | 84.75 | 75.77 | 63.60 | 69.15 | 60.80 | 60.97 | 60.88 |
| *Solanum lycopersicum* | 79.64 | 66.54 | 72.50 | 46.12 | 37.27 | 41.23 | 46.12 | 34.58 | 39.52 |
| **Average** | 85.09 | 80.74 | 82.69 | 66.87 | 54.79 | 59.62 | 53.63 | 51.42 | 51.30 |

Supplemental Table S10: Sensitivity, precision, and F1-score of the AUGUSTUS predictions made as part of the BRAKER3 pipeline. The results correspond to Supplemental Table S4 – proteins from the same order as the target species were excluded.

| | Exon | | | Gene | | | Transcript | | |
|---|---|---|---|---|---|---|---|---|---|
| | Sn | Prec | F1 | Sn | Prec | F1 | Sn | Prec | F1 |
| Funannotate | 73.20 | 74.40 | 73.79 | 47.11 | 44.69 | 45.87 | 35.75 | 44.70 | 39.73 |
| Funannotate updated | 73.20 | 74.40 | 73.79 | 47.11 | 44.69 | 45.87 | 35.75 | 44.70 | 39.73 |
| Funannotate –repeats2evm | 71.12 | 77.06 | 73.97 | 45.49 | 45.64 | 45.56 | 34.45 | 45.64 | 39.27 |
| Funannotate –repeats2evm updated | 73.20 | 74.40 | 73.79 | 47.11 | 44.69 | 45.87 | 35.75 | 44.70 | 39.73 |

Supplemental Table S11: Average sensitivity, precision, and F1-score for four different predictions generated by Funannotate using the **close relatives included** protein databases for the same species as listed in Supplemental Table S8. The prediction step of Funannotate was run with and without the option to pass gene predictions of repetitive regions to EVidenceModeler (`--repeats2evm`). The resulting predictions were both post-processed using Funannotate's update protocol, which updates the predicted gene models using RNA-seq data.

| | *Arabidopsis thaliana* | *Bombus terrestris* | *Caenorhabditis elegans* | *Danio rerio* |
|---|---|---|---|---|
| BRAKER1 | 01:23 | 04:16 | 01:36 | 10:28 |
| BRAKER2 | 06:39 | 06:27 | 04:57 | 19:47 |
| GeneMark-ETP | 04:05 | 02:56 | 03:36 | 09:54 |
| BRAKER3 | 05:37 | 14:45 | 06:46 | 27:08 |
| | *Drosophila melanogaster* | *Gallus gallus* | *Medicago truncatula* | *Mus musculus* |
| BRAKER1 | 01:55 | 10:12 | 03:20 | 19:41 |
| BRAKER2 | 05:28 | 14:10 | 10:59 | 30:24 |
| GeneMark-ETP | 02:37 | 12:11 | 07:55 | 22:13 |
| BRAKER3 | 06:50 | 42:44 | 13:14 | 64:16 |
| | *Parasteatoda tepidariorum* | *Populus trichocarpa* | *Solanum lycopersicum* | **Average** |
| BRAKER1 | 14:47 | 04:41 | 03:47 | 06:55 |
| BRAKER2 | 23:16 | 08:52 | 09:19 | 12:45 |
| GeneMark-ETP | 10:38 | 07:51 | 06:37 | 08:14 |
| BRAKER3 | 35:35 | 11:41 | 12:06 | 21:53 |

Supplemental Table S12: Runtime of BRAKER1, BRAKER2, GeneMark-ETP, and BRAKER3 for all test species. The runtime is written as hours and minutes. The hardware is described in the caption of Supplemental Figure S6.

| Species | Runtime (h:m) |
|---|---|
| Bombus terrestris | 4:00 |
| Drosophila melanogaster | 2:06 |
| Gallus gallus | 13:54 |
| Medicago truncatula | 2:48 |
| Populus trichocarpa | 3:36 |

Supplemental Table S13: Runtime of MAKER2 with *close relatives included* database in MPI mode using LINUX nodes with 96 cores available through the Azure cloud. Training of *ab initio* gene finders and transcriptome assembly are not included in these figures.

| | *Arabidopsis thaliana* | *Bombus terrestris* | *Caenorhabditis elegans* | *Drosophila melanogaster* |
|---|---|---|---|---|
| Funannotate | 05:06 | 04:59 | 02:02 | 02:52 |
| BRAKER3 | 05:46 | 14:38 | 04:13 | 06:25 |
| | *Gallus gallus* | *Medicago truncatula* | *Populus trichocarpa* | *Solanum lycopersicum* |
| Funannotate | 41:07 | 07:11 | 07:03 | 09:28 |
| BRAKER3 | 27:58 | 07:54 | 08:19 | 08:25 |
| | **Average** | | | |
| Funannotate | 09:59 | | | |
| BRAKER3 | 10:27 | | | |

Supplemental Table S14: Runtime of Funannotate and BRAKER3 for the experiments of Supplemental Table S8 (*close relatives included*). The runtime is written as hours and minutes. The hardware is described in the caption of Supplemental Figure S6.

| Species | SRA ID | #spots | #bases | Date |
|---|---|---|---|---|
| *A. tha.* | SRR8714016 | 26,518,474 | 8,008,579,148 | 2020-02-27 |
| | SRR8759751 | 21,388,481 | 6,458,011,735 | 2019-04-01 |
| | SRR4010853 | 25,658,415 | 6,465,920,580 | 2017-05-10 |
| | SRR7289569 | 9,222,308 | 1,391,511,444 | 2018-12-06 |
| | SRR12547664 | 874,772 | 66,482,672 | 2021-04-07 |
| | SRR12076896 | 21,857,417 | 6,557,225,100 | 2021-05-24 |
| *B. ter.* | SRR5125126 | 28,930,495 | 5,207,489,100 | 2017-10-30 |
| | SRR5125123 | 25,185,030 | 4,533,305,400 | 2017-10-30 |
| | SRR16931591 | 25,056,341 | 7,516,902,300 | 2021-11-12 |
| | SRR5125133 | 26,059,038 | 4,690,626,840 | 2017-10-30 |
| | SRR5125134 | 23,745,975 | 4,274,275,500 | 2017-10-30 |
| | SRR8085469 | 32,768,117 | 6,553,623,400 | 2019-03-05 |
| *C. ele.* | SRR7446944 | 44,853,824 | 8,970,764,800 | 2020-06-01 |
| | ERR2756716 | 4,785,656 | 1,445,268,112 | 2018-08-29 |
| | SRR6815567 | 5,901,194 | 890,097,086 | 2018-03-08 |
| | SRR6474814 | 34,199,521 | 8,618,279,292 | 2018-01-16 |
| | SRR10238291 | 47,652,843 | 4,955,895,672 | 2019-11-07 |
| *D. rer.* | SRR3179613 | 9,994,443 | 2,018,877,486 | 2017-02-10 |
| | ERR1857957 | 2,536,494 | 380,474,100 | 2017-03-03 |
| | SRR9159941 | 19,527,035 | 5,858,110,500 | 2019-12-01 |
| | SRR9159937 | 37,156,875 | 11,147,062,500 | 2019-12-01 |
| | ERR958944 | 10,490,607 | 1,363,778,910 | 2015-07-17 |
| | SRR8106574 | 1,281,189 | 186,371,148 | 2020-01-20 |
| *D. mel.* | SRR19416937 | 21,377,196 | 6,413,158,800 | 2022-05-30 |
| | SRR19416947 | 22,611,252 | 6,783,375,600 | 2022-05-30 |
| | SRR19416944 | 17,754,393 | 5,326,317,900 | 2022-05-30 |
| | SRR19446462 | 40,819,492 | 4,163,588,184 | 2022-05-30 |
| | SRR19416948 | 21,257,788 | 6,377,336,400 | 2022-05-30 |
| *M. tru.* | SRR3735569 | 13,021,000 | 2,630,242,000 | 2016-07-02 |
| | SRR2016009 | 4,289,979 | 428,997,900 | 2016-05-12 |
| | SRR10416790 | 9,757,388 | 2,927,216,400 | 2019-11-23 |
| | SRR3726824 | 44,634,145 | 13,390,243,500 | 2016-06-28 |
| | SRR2015998 | 7,916,226 | 791,622,600 | 2016-05-12 |
| | SRR15462191 | 21,367,770 | 6,410,331,000 | 2021-08-15 |
| *P. tep.* | SRR5458595 | 94,367,928 | 18,873,585,600 | 2017-06-14 |
| | SRR1824488 | 33,099,205 | 6,686,039,410 | 2015-03-03 |
| | SRR12687629 | 41,410,278 | 12,423,083,400 | 2021-06-22 |
| | SRR8755634 | 45,919,505 | 8,852,599,087 | 2019-03-20 |
| | SRR5602551 | 9,600,853 | 2,400,213,250 | 2017-10-16 |
| | SRR1824489 | 60,177,256 | 12,155,805,712 | 2015-03-03 |
| *P. tri.* | SRR3019959 | 32,156,743 | 4,591,697,381 | 2016-01-19 |
| | SRR3019957 | 44,863,324 | 6,484,994,933 | 2016-01-19 |
| | SRR3019251 | 29,773,370 | 4,305,817,693 | 2016-01-19 |
| | SRR3019585 | 36,843,662 | 5,279,588,065 | 2016-01-19 |
| | SRR3019304 | 41,358,002 | 5,970,634,746 | 2016-01-19 |
| | SRR12671667 | 17,429,154 | 5,263,604,508 | 2020-09-18 |
| *G. gal.* | SRR5340686 | 27,956,970 | 2,851,610,940 | 2017-03-15 |
| | ERR2113192 | 20,988,313 | 3,148,246,950 | 2017-09-07 |
| | ERR2113173 | 21,562,175 | 3,234,326,250 | 2017-09-07 |
| | SRR5190436 | 23,938,529 | 4,787,705,800 | 2017-09-06 |
| | SRR1822373 | 48,874,648 | 7,819,943,680 | 2015-05-05 |
| | SRR5437696 | 35,378,254 | 5,306,738,100 | 2017-09-18 |
| *M. mus.* | SRR5197958 | 10,577,301 | 3,184,479,151 | 2017-08-31 |
| | SRR3094250 | 1,568,252 | 316,786,904 | 2016-11-14 |
| | SRR6067921 | 5,886,768 | 888,143,922 | 2018-08-03 |
| | SRR10115888 | 44,307,998 | 8,861,599,600 | 2020-10-02 |
| | ERR3005082 | 8,598 | 1,289,700 | 2019-01-02 |
| | SRR9202226 | 5,355,073 | 546,217,446 | 2019-08-07 |

Supplemental Table S15: This table lists all RNA-seq libraries used for each species in all experiments. It includes the ID of the library at the Sequence Read Archive, the number of spots, number of bases, and publication date.

| Species | AUGUSTUS | GeneMark.hmm | SNAP |
|---|---|---|---|
| *A. thaliana* | arabidopsis* | from GeneMark-ES run | A.thaliana.hmm* |
| *B. terrestris* | bombus_terrestris2* | from GeneMark-EP+ run | trained with genes from ref. annotation |
| *C. elegans* | c_elegans_trsk* | from GeneMark-ES run | Ce.hmm* |
| *D. melanogaster* | fly* | from GeneMark-ES run | fly* |
| *G. gallus* | chicken* | medium GC model from GeneMark-ETP run | trained with genes from ref. annotation |
| *M. truncatula* | trained with genes from ref. annotation | from GeneMark-EP+ run | trained with genes from ref. annotation |
| *P. trichocarpa* | trained with genes from ref. annotation | from GeneMark-EP+ run | trained with genes from ref. annotation |
| *S. lycopersicum* | tomato* | from GeneMark-EP+ run | trained with genes from ref. annotation |

Supplemental Table S16: Configuration of AUGUSTUS, GeneMark.hmm and SNAP gene models using parameter from the AUGUSTUS and the SNAP distribution packages (marked with *) or with manually training using a training gene set.

#### 1.3 Supplementary Methods

##### Running BRAKER

BRAKER v3.0.2 was installed from GitHub (`https://github.com/Gaius-Augustus/BRAKER`) and the BRAKER pipelines were run as follows. The GeneMark-ETP results were taken from the BRAKER3 runs:

BRAKER1:

```
braker.pl --genome=genome.softmasked.fasta --threads=48 \
    --rnaseq_sets_dirs=/location/of/local/RNA_Seq/Files \
    --rnaseq_sets_ids=RNA_Seq_IDS
```

BRAKER2:

```
braker.pl --genome=genome.softmasked.fasta --prot_seq=proteins.fa \
     --threads=48
```

BRAKER3:

```
braker.pl --genome=genome.softmasked.fasta --prot_seq=proteins.fa \
    --rnaseq_sets_dirs=/location/of/local/RNA_Seq/Files \
    --rnaseq_sets_ids=RNA_Seq_IDS --threads=48
```

##### Running Funannotate

Funannotate v1.8.14 was installed using a Singularity container as follows:

```
# only once, to get the singularity container
singularity pull docker://nextgenusfs/funannotate

export GENEMARK_PATH=/path/to/GeneMark-ES-ET-EP_v4.71_lic

species="name of species"
buscoSeedSpecies="name of seed species"
buscodb="name of busco db"
genome="/path/to/genome.fasta.masked"
proteins="/path/to/proteins.fa"
bamfile="/path/to/bamfile.bam"

# calculateGenomeSizeFromFasta.pl adds up the length of all sequences in a fasta
genomeSize=$(perl ~/calculateGenomeSizeFromFasta.pl $genome)
maxIntronLen_f=$(echo "3.6 * sqrt($genomeSize)" | bc -l)
maxIntronLen=$(printf "%.0f" "$maxIntronLen_f")

singularity run funannotate_latest.sif funannotate sort \
    --input $genome \
    --out genome.sorted.fa \
    --simplify
```

We produced four different results for each species, using the option `--repeats2evm` during the prediction step and a recommended update step:

```
mkdir -p fun tmp
# run prediction
singularity run funannotate_latest.sif funannotate predict \
    --input genome.sorted.fa --out fun --species $species \
    --busco_seed_species $buscoSeedSpecies --busco_db $buscodb \
    --organism other --protein_evidence $proteins \
    --rna_bam $bamfile --max_intronlen $maxIntronLen \
    --cpus 72 --tmpdir tmp --no-progress \
    [--repeats2evm]
```

```
# run update step
singularity run funannotate_latest.sif funannotate update \
  --input fun/ --left $readL --right $readR \
  --max_intronlen $maxIntronLen --species $species \
  --memory 50G --cpus 72 --no-progress
```

##### Running FINDER

FINDER v1.1.0 was installed from GitHub (`https://github.com/sagnikbanerjee15/Finder`) and run as follows:

```
run_finder --protein proteins.fa --framework singularity --output_directory finder --cpu 48 \
    --metadatafile metadata.csv --genome genome.softmasked.fasta --genemark_license gm_key \
    --genemark_path /location/of/GeneMark-ES/ET/EP  --organism_model {VERT,INV,PLANTS,FUNGI}
```

The information in the `metadata.csv` files was manually generated for each species and includes details about the RNA-seq libraries. The files consist of the following fields:

```
BioProject, SRA Accession, Tissues, Description, Date, Read Length (bp),
Ended, RNA Seq, process, Location
```

##### Running MAKER2

We ran MAKER2 version 3.01.04 using AUGUSTUS version 3.5.0, GeneMark.hmm version 3.68, SNAP version 2.59.5, Exonerate version 2.2.0, BLAST version 2.14.0, Tandem Repeats Finder version 4.09.

We provided MAKER with a GFF file containing the coordinates of repeats, as well as their predicted repeat types from RepeatMasker. To ensure compatibility with MAKER, we reformatted the RepeatMasker output to a specific GFF format using the following script from `https://github.com/gatech-genemark/BRAKER2-exp.git` and command line:

```
rmasker_out2maker_gff.pl < genome.fasta.out > rmasker4maker.gff
```

This script is available from the repository `https://github.com/gatech-genemark/GeneMark-ETP-exp`.

The default MAKER2 configuration was adjusted with following settings:

```
genome=genome.fasta
est=transcriptome.fasta
protein=proteindb.fasta
model_org=   #empty
rm_gff=repeatmasker.gff
snaphmm=snap.model
gmhmm=genemark.mod
augustus_species=model_name
est2genome=1
protein2genome=1
keep_preds=0
```

Here, `transcriptome.fasta` contained the same transcriptome assemblies that were used with BRAKER3 and constructed with HISAT2 and StringTie2. MAKER2 was then run with:

```
mpiexec.mpich -n 96  maker
```

##### Preparing protein data

OrthoDB v.11 was partitioned into proteins of species from the clade *Arthropoda*, *Metazoa*, *Vertebrata*, and *Viridiplantae*. The partitioning is available from `https://bioinf.uni-greifswald.de/bioinf/partitioned_odb11/`. Subsequently, two protein sets for each species from Suppl. Table 1 were generated, excluding either only the target species (species-excluded) or all species of the same taxonomic order (order-excluded).

These sets were prepared using the orthodb-clades pipeline, downloaded from GitHub (`https://github.com/tomasbruna/orthodb-clades`).

```
snakemake --cores 48
```

##### Accuracy evaluation

The performance measurements were computed using scripts from the BRAKER suite:

```
compute_accuracies.sh ref_annot.gtf pseudo.gff3 gene_set.gtf gene trans cds
```

##### Measuring expression

Expression levels were estimated with `kallisto` version 0.50.0 (Bray et al., 2016). For each species, the mRNA sequences of the respective reference annotation were quantified using a pool of RNA-seq libraries. More specifically, all paired libraries that were used for annotation were pooled and TPM (transcript per million) values estimated with the following command line.

```
kallisto quant -i kallisto.idx -o work/ -t 8 --verbose input/*.fastq
```

Subsequently, the transcripts for each species were partitioned in three expression *terciles*: the first with lowest TPM value, the second with medium TPM value and the third with highest TPM value.
